# Single chromosome gains can function as metastasis suppressors and metastasis promoters in colon cancer

**DOI:** 10.1101/590547

**Authors:** Anand Vasudevan, Prasamit S. Baruah, Joan C. Smith, Zihua Wang, Nicole M. Sayles, Peter Andrews, Jude Kendall, Justin E. Leu, Narendra Kumar Chunduri, Dan Levy, Michael Wigler, Zuzana Storchová, Jason M. Sheltzer

## Abstract

Most human tumors display chromosome-scale copy number alterations, and high levels of aneuploidy are frequently associated with advanced disease and poor patient prognosis. To examine the relationship between aneuploidy and cancer progression, we generated and analyzed a series of congenic human cell lines that harbor single extra chromosomes. We find that different aneuploidies can have distinct effects on invasive behavior: across 13 different cell lines, 12 trisomies suppressed invasiveness or were largely neutral, while a single trisomy increased metastatic behavior by triggering a partial epithelial-mesenchymal transition. In contrast, chromosomal instability, which can lead to the development of aneuploidy, uniformly suppressed cellular invasion. By analyzing genomic copy number and survival data from 10,133 cancer patients, we demonstrate that specific aneuploidies are associated with distinct clinical outcomes, and the acquisition of certain aneuploidies is in fact linked with a favorable prognosis. Thus, aneuploidy is not a uniform driver of malignancy, and different chromosome copy number changes can uniquely influence tumor progression. At the same time, the gain of a single chromosome is capable of inducing a profound cell state transition, underscoring how genomic plasticity can engender phenotypic plasticity and lead to the acquisition of enhanced metastatic properties.

## Introduction

Whole-chromosome aneuploidy is a nearly-ubiquitous feature of human tumors, though its role in malignancy remains poorly understood (Sheltzer and Amon 2011; Knouse et al. 2017; Chunduri and Storchová 2019). Aneuploidy alters the dosage of hundreds or thousands of genes at once, causing proportional changes in the expression of most transcripts on an affected chromosome (Williams et al. 2008; Sheltzer et al. 2012; Stingele et al. 2012; Dürrbaum et al. 2014). These dosage imbalances have pleiotropic effects on cell physiology, and can impair metabolism, protein homeostasis, and the maintenance of genomic stability (Williams et al. 2008; Sheltzer et al. 2011; Donnelly et al. 2014; Passerini et al. 2016). Despite its prevalence in cancer, under most conditions aneuploidy inhibits, rather than promotes, cell proliferation, and many single-chromosome gains can suppress tumorigenesis (Williams et al. 2008; Sheltzer et al. 2017). At the same time, the phenotypic alterations caused by aneuploidy may lead to improved fitness in certain environments (Pavelka et al. 2010; Rutledge et al. 2016), and aneuploidy can serve as a mechanism by which cells increase the dosage of growth-promoting oncogenes (Davoli et al. 2013). Thus, aneuploidy can have multifaceted and sometimes opposing roles during tumor development.

Aneuploidy can arise as a result of a cellular condition called chromosomal instability, or CIN (Holland and Cleveland 2009). Under normal conditions, the cell’s spindle assembly checkpoint (SAC) maintains a metaphase arrest until all chromosomes are correctly aligned and under tension from the mitotic spindle (Lara-Gonzalez et al. 2012). The kinase Mps1 is the master regulator of this checkpoint, and it phosphorylates multiple proteins capable of measuring the tension status of each chromosome (Pachis and Kops 2018). If the SAC is compromised, cells will undergo anaphase prematurely, leading to the random missegregation of chromosomes and the development of aneuploidy.

Clinically, cancer aneuploidy has been recognized as a common indicator of poor patient prognosis (Merkel and McGuire 1990). The overall degree of aneuploidy tends to increase during tumor progression, and aneuploid tumors from several tissues are correlated with decreased overall survival compared to tumors with near-diploid karyotypes (Sheltzer 2013; Frankfurt et al. 1985; Friedlander et al. 1984; Choma et al. 2001; Kheir et al. 1988; Kallioniemi et al. 1987). However, most of these analyses have relied on technologies that are only capable of detecting gross deviations from the euploid state (e.g., DNA flow cytometry). Modern array-based and sequencing-based methods can be used to determine the complete set of karyotypic alterations found in a patient’s tumor, but the prognostic importance of most specific aneuploidies is not known. Additionally, the reasons underlying the close link between aneuploidy and outcome remain contested.

As the vast majority of cancer deaths result from metastasis, it is possible that aneuploidy plays a role in the metastatic cascade (Mehlen and Puisieux 2006). During metastasis, cancer cells must migrate out of their local environment, intravasate into the vascular system, travel to a distant organ, and re-establish tumor growth at a secondary site (Pantel and Brakenhoff 2004). In epithelial cancers, this process has been associated with the co-option of a developmental program called the epithelial-mesenchymal transition (EMT) (Heerboth et al. 2015). During EMT, cancer cells silence the expression of the cell-adhesion proteins typically found in epithelial tissue and activate a set of mesenchymal genes that drive cell movement and invasion. While the expression of an EMT gene signature has been correlated with poor prognosis (Taube et al. 2010), further experiments have challenged the notion that the EMT is either necessary or sufficient for tumor metastasis (Zheng et al. 2015; Fischer et al. 2015). Related programs, including a partial-EMT, in which cells fail to completely adopt a mesenchymal phenotype, have also been proposed to play a role in cancer cell dissemination (Jolly et al. 2017; Pastushenko and Blanpain 2019).

Alternately, it is possible that aneuploidy does not directly affect metastasis, and instead arises in aggressive cancers as a by-product of tumor evolution. For instance, mutations in the tumor suppressor *TP53* are associated both with poor prognosis and with the acquisition of aneuploidy (Thompson and Compton 2010; López-García et al. 2017; Smith and Sheltzer 2018), and the increased levels of aneuploidy in late-stage malignancies could simply be a consequence of the loss of p53 function. Finally, a recent study has suggested that CIN, rather than aneuploidy *per se*, is a key driver of cancer metastasis (Bakhoum et al. 2018). According to this hypothesis, lagging chromosomes caused by CIN become trapped outside of the nucleus, forming micronuclei, and are recognized by the cGAS/STING cytosolic DNA sensing pathway. STING triggers the activation of non-canonical NFκB signaling, leading to an EMT and increased cellular invasiveness (Bakhoum et al. 2018). Thus, in this model, the observed correlation between aneuploidy and metastasis may occur as a byproduct of the relationship between aneuploidy and CIN.

Our ability to understand the connections between CIN, aneuploidy, and metastasis has been hampered by a lack of controlled systems in which to study these processes. The aneuploidy that naturally develops during tumorigenesis is often triggered or accompanied by additional mutations, compromising our ability to determine causal relationships. In order to overcome these limitations, we and others have developed multiple approaches to generate human cell lines with specific whole-chromosome aneuploidies (Williams et al. 2008; Stingele et al. 2012; Sheltzer et al. 2017). By altering chromosome copy number in an otherwise-identical background, we can isolate and study the effects of aneuploidy, absent any other secondary alterations. Similarly, while the causes of CIN in human tumors remain debated, chromosomal instability can be generated *in vitro* using small-molecule inhibitors of the SAC kinase Mps1, allowing us to isolate the specific consequences of CIN (Santaguida et al. 2010, 2015; Sheltzer et al. 2017). Using these systems, we set out to investigate how aneuploidy and CIN affect metastatic behavior in cancer.

## Results

### Single-chromosome gains can either promote or suppress metastatic behavior

Using microcell-mediated chromosome transfer (MMCT), we generated derivatives of the human colon cancer cell line HCT116 that harbored single extra chromosomes (Stingele et al. 2012; Domingues et al. 2017). The parental cell line exhibits a near-diploid karyotype, with partial gains of chromosomes 8, 10, 16, and 17. We successfully derived clones of this line that harbored additional trisomies of one of six different chromosomes (Chr. 3, 5, 8, 13, 18, or 21). We then confirmed the presence of the extra chromosome and the absence of any secondary aneuploidies in each clone using Short Multiply Aggregated Sequence Homologies (SMASH) sequencing (Figure S1A)(Wang et al. 2016).

To test how these alterations affected metastatic behavior, we applied an *in vitro* invasion assay, in which cells were challenged by a chemotactic gradient to invade through a basement membrane (Figure 1A). In the near-diploid parental HCT116 line, about 20 cells per field were able to successfully invade through the Matrigel layer (Figure 1B). Gaining an extra copy of Chr. 3 or Chr. 21 did not affect the rate of invasion, while gaining an extra copy of Chr. 13 or 18 significantly decreased invasion. However, in four independent clones trisomic for Chr. 5, cellular invasion significantly increased to nearly 100 cells per field (Figure 1B-C).

**Figure 1.**
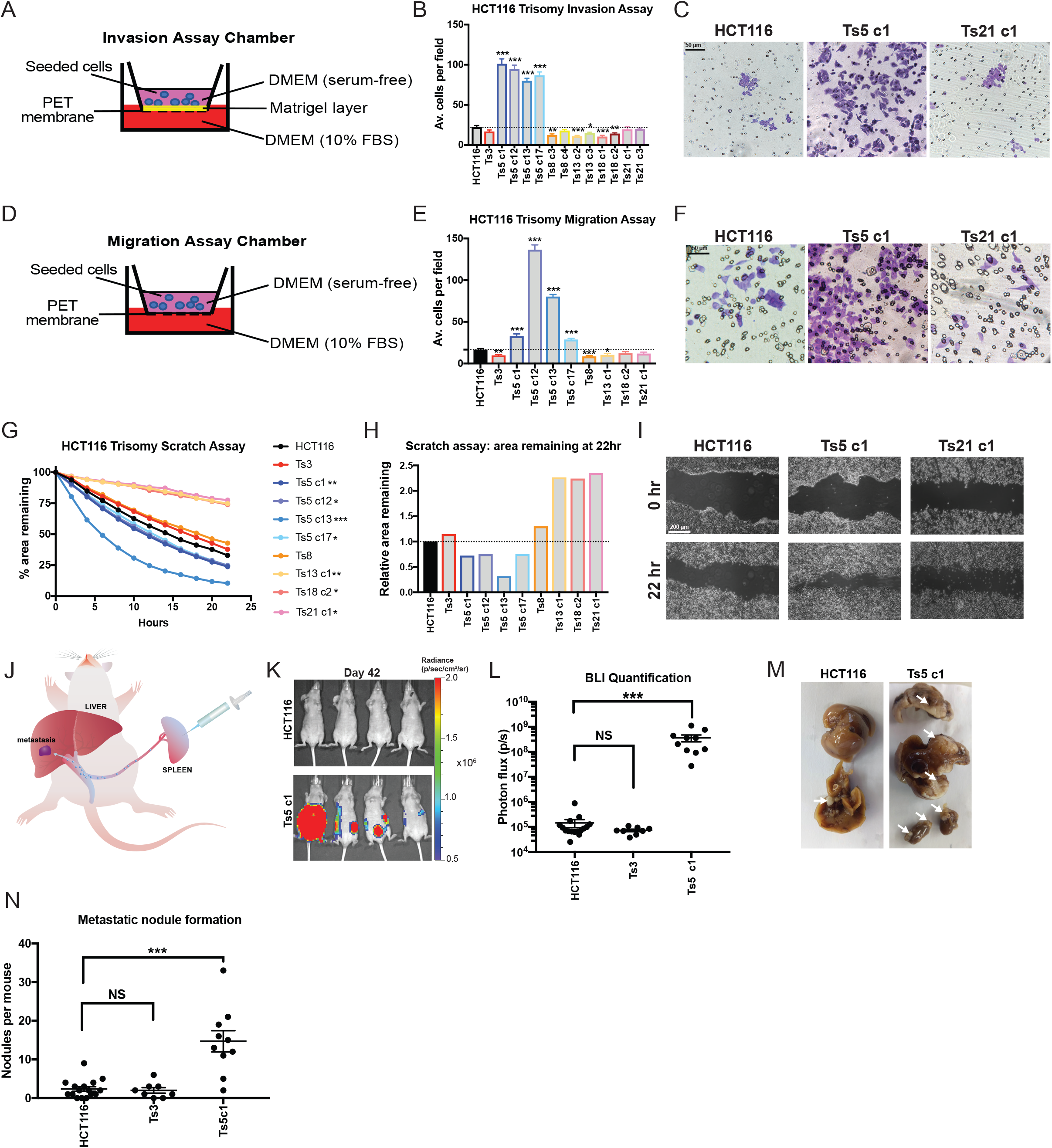
Gaining chromosome 5 increase invasion, migration, and motility in colon cancer cells. (A) Schematic diagram of the invasion assay. Cells are seeded in the upper chamber in serum-free media while the bottom chamber contains media supplemented with 10% FBS. (B) Quantification of the average number of cells per field that were able to cross the membrane in the invasion assay. Averages represent three independent trials in which 15-20 fields were counted. (C) Representative images of HCT116, HCT116 Ts5 c1, and HCT116 Ts21 c1 invasion stained with crystal violet. (D) Schematic diagram of migration assay. The setup is similar to the invasion assay except that the membrane does not have a matrigel layer. (E) Quantification of the average number of cells per field that were able to cross the membrane in the migration assay. Averages represent three independent trials in which 15-20 fields were counted. (F) Representative images of HCT116, HCT116 Ts5 c1, and HCT116 Ts21 c1 migration stained with crystal violet. (G) Quantification of cell motility in a scratch assay. The percent area remaining between two monolayers separated by a pipette tip-induced scratch was monitored for 22 hours. Minimum of 2 trials per cell line. (H) The ratio of the area remaining at 22 hours after the scratch is plotted relative to the HCT116 parental cell line. A ratio less than 1 indicates faster scratch closure relative to wild-type, while a ratio greater than 1 indicates slower scratch closure. (I) Representative images of the scratch closure assay immediately after the scratch (0h) and at the end of the assay (22h) for HCT116, HCT116 Ts5 c1, and HCT116 Ts21 c1. Bars represent mean ± SEM. * p<0.05; ** p<0.005 *** p<0.0005. Scale bar, 50 and 200 µm (J) A schematic of the colon cancer metastasis assay. Luciferase-expressing cells are injected into the mouse spleen, and then metastatic nodule formation in other organs is quantified. (K) Luciferase imaging of mice injected with HCT116 or HCT116 Ts5 cells 42 days post-injection. (L) Bioluminescence imaging quantification of the luciferase signal in mice injected with HCT116, HCT116 Ts3, and HCT116 Ts5 cells. (M) Images of metastatic nodule formation from mice euthanized 42 days post-injection. White arrows indicate metastases. (N) Quantification of the number of nodules per mouse (HCT116 n = 16, HCT116 Ts3 n = 8, HCT116 Ts5 c1 n = 10). Bars represent mean ± SEM. Unpaired t-test * p<0.05; ** p<0.005 *** p<0.0005. Scale bar, 50 and 200 µm.

To further explore the effects of aneuploidy on metastasis-related phenotypes, we performed two additional assays in each trisomic cell line. First, we conducted a migration assay, in which cells were challenged by a chemotactic gradient to move through an 8 µm pore (Figure 1D-F). Secondly, we performed a scratch-repair assay, in which we used live-cell imaging to follow the rate at which each cell line was capable of closing a cell-free gap (Figure 1G-I). From these assays, certain patterns emerged: most trisomic lines were either neutral or displayed decreased invasiveness in each experiment. HCT116 Ts13, in particular, exhibited significantly lower rates of cellular invasion and migration, and moved nearly twice as slowly as the parental line in the scratch closure assay. In contrast, four of four tested clones that harbored an extra copy of chromosome 5 displayed increased metastatic behavior in each assay. We noted some variability between independent Ts5 clones (e.g., compare Ts5 c1 and Ts5 c12, Figure 1E), but the overall patterns were consistent across this trisomy. In total, these assays demonstrate a complicated relationship between single-chromosome aneuploidies and metastatic phenotypes: specific aneuploidies may either promote, suppress, or fail to affect different metastasis-related processes, and gaining chromosome 5 in particular has a consistently strong effect on invasiveness in this colon cancer cell line.

Next, to test whether the patterns that we observed *in vitro* translated to changes in metastatic behavior *in vivo*, we injected luciferase-expressing HCT116, HCT116 Ts3, or HCT116 Ts5 cells into the spleens of nude mice and then quantified the formation of metastatic nodules on the liver and other organs (Figure 1J)(Morikawa et al. 1988). Under these conditions, the wild-type cell line was very weakly metastatic, as evidenced by the lack of luciferase signal and the scarcity of metastatic nodules that had formed by six weeks post-injection (Figure 1K-M). HCT116 Ts3 cells were also poorly metastatic, forming a similar number of nodules as the parental line. (Figure 1K). However, cells trisomic for chromosome 5 exhibited substantial metastatic activity and formed on average 15 nodules per mouse, significantly more than the parental line (P < .0001, Student’s t-test). We conclude that the amplification of chromosome 5 in HCT116 cells increases invasive behavior both *in vitro* and *in vivo*.

We then set out to investigate the impact of aneuploidy in other human cell lines. Using MMCT, we generated TERT-transformed retinal pigment epithelial (RPE1) cell lines trisomic for Chr. 3, 5, 7, or 21 (Stingele et al. 2012; Domingues et al. 2017; Dürrbaum et al. 2018). Though these cells are non-cancerous, they display several hallmarks of malignant growth (activation of RAS-MAPK signaling, rapid cell division, immortality, and invasive behavior) and they provide an independent genetic background to interrogate the effects of aneuploidy (Di Nicolantonio et al. 2008). Additionally, human developmental trisomies like Down syndrome (Ts21) are commonly associated with neural tube defects and other CNS abnormalities that could potentially result from impaired migration, and we sought to uncover whether single chromosome gains could also affect invasion and motility in non-tumor derived cells (Hume et al. 1996; Becker et al. 1991). We therefore tested the behavior of the trisomic RPE1 cell lines in the assays described above. RPE1 Ts3, Ts5, Ts7, and Ts21 were found to suppress metastatic behavior, and Ts3 had a particularly strong inhibitory effect (Figure S2A-E). While Ts5 increased invasiveness in HCT116 cells, in RPE1 this extra chromosome consistently blocked invasive behavior. This suggests that unique interactions between an aneuploidy and the cell’s genetic background determine whether a specific chromosome gain will have pro-metastatic or anti-metastatic consequences. Interestingly, gaining chromosome 21 strongly decreased monolayer closure in the scratch-repair assay, suggesting that some of the CNS defects observed in Down syndrome could result from Ts21-induced dysregulation of cell motility (Figure S2D-E).

Finally, to ensure that our results were not a consequence of the MMCT protocol used to generate these aneuploidies, we applied an alternate methodology to derive trisomic cells. We treated near-diploid DLD1 colon cancer cells with AZ3146, a small-molecule inhibitor of the SAC kinase Mps1, and isolated single cell-derived clones. The DLD1 parental cell line harbors partial or focal gains on chromosomes 2, 11, and 16, and we successfully derived clones with additional whole-chromosome trisomies of either Chr. 4, 6, or 11 (Figure S1C). We subjected these clones to each metastasis assay, and we found that these trisomies were either neutral or suppressed invasive behavior (Figure S2F-J). In total, our results indicate that aneuploidy can have complex effects on metastasis-related phenotypes. Single-chromosome aneuploidies can be sufficient to either promote or suppress invasiveness, depending on the cell’s genetic background and the identity of the aneuploid chromosome.

### Trisomy 5 promotes invasive behavior by causing a partial epithelial-mesenchymal transition and upregulating matrix metalloproteinases

We next sought to investigate how gaining chromosome 5 in HCT116 causes an increase in invasiveness. As metastasis is commonly associated with an EMT, we assessed the expression of canonical markers of epithelial cell identity in each clone of HCT116 trisomy. We found that wild-type HCT116 cells and HCT116 derivatives trisomic for Chr. 3, 8, 13, 18, or 21 expressed high levels of the epithelial adhesion genes EpCAM, E-cadherin, and Claudin 7 (Figure 2A). In contrast, EpCAM, E-cadherin, and Claudin 7 were silenced in nine out of ten clones trisomic for chromosome 5, suggesting that this aneuploidy caused a loss of epithelial cell identity. To test whether Ts5 converted cells to a mesenchymal state, we next assessed the expression of the mesenchymal markers Fibronectin, N-cadherin, and Vimentin. We found that these genes were not expressed in any trisomic derivative of HCT116, including Ts5, indicating that this aneuploidy induces a “partial” EMT phenotype (Figure S3).

**Figure 2.**
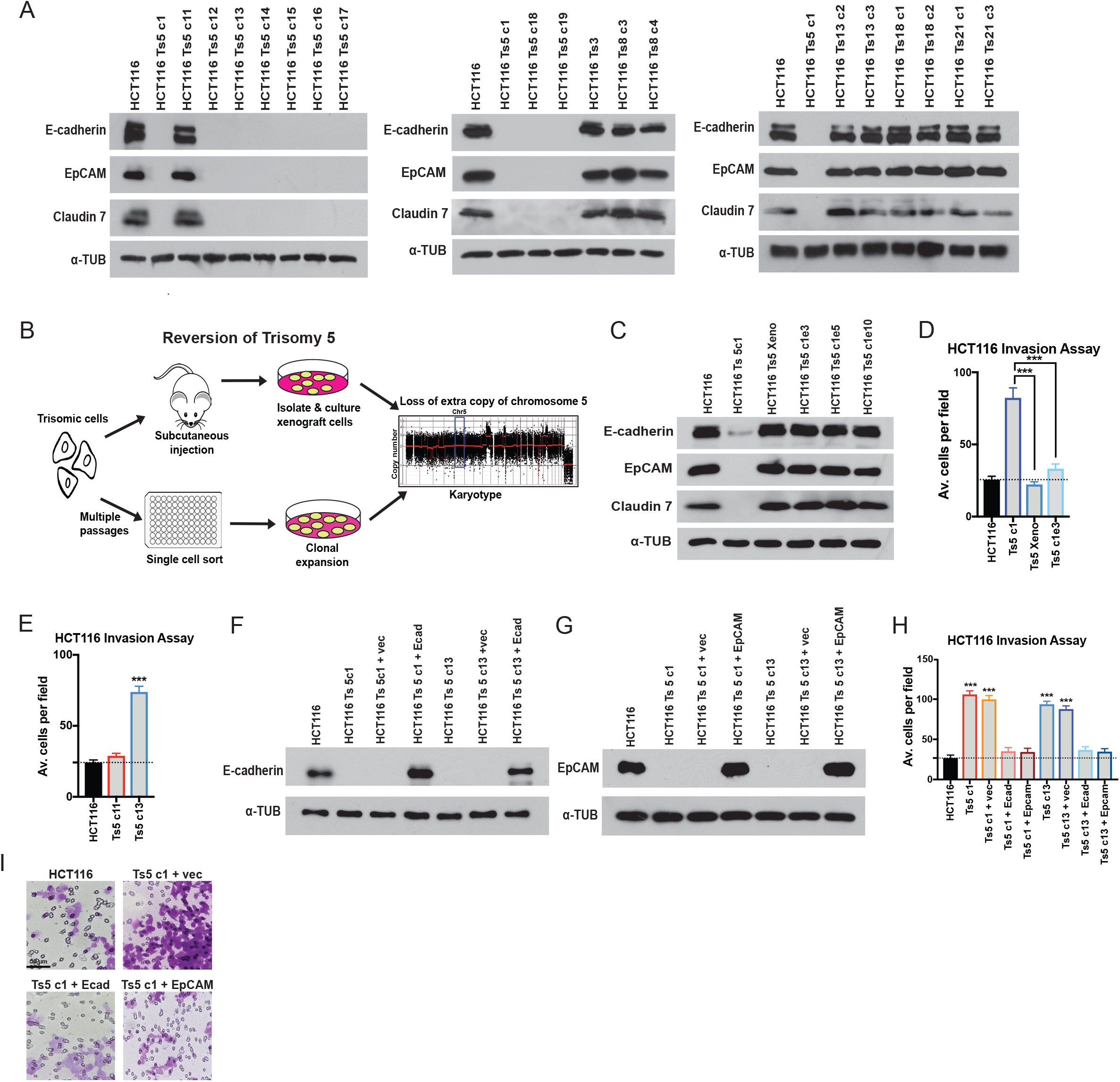
Gaining chromosome 5 induces a cell state transition in colon cancer cells. (A) Western blot analysis of epithelial marker expression in HCT116 trisomies. Alpha-tubulin was used as a loading control. (B) Schematic of two strategies to select for HCT116 Ts5 chromosome-loss revertants. (C) Western blot analysis of epithelial marker expression in trisomy 5 cells that lost the extra copy of chromosome 5. “Xeno”: cells isolated after xenograft growth. Ts5 c1 e3, e5, e10: clones isolated from high-passage cells. (D-E) Quantification of the average number of cells per field that were able to cross the membrane in the invasion assay. Averages represent three independent trials in which 15-20 fields were counted. (F-G) Verification of E-cadherin and EpCAM overexpression in two HCT116 Ts5 clones transfected with the indicated plasmids. (H) Quantification of the average number of cells per field that were able to cross the membrane in the invasion assay. Averages represent three independent trials in which 15-20 fields were counted. (I) Representative images of invasion in the indicated cell lines. Bars represent mean ± SEM. Unpaired t-test *** p<0.0005. Scale bar, 50 µm

**Figure 3.**
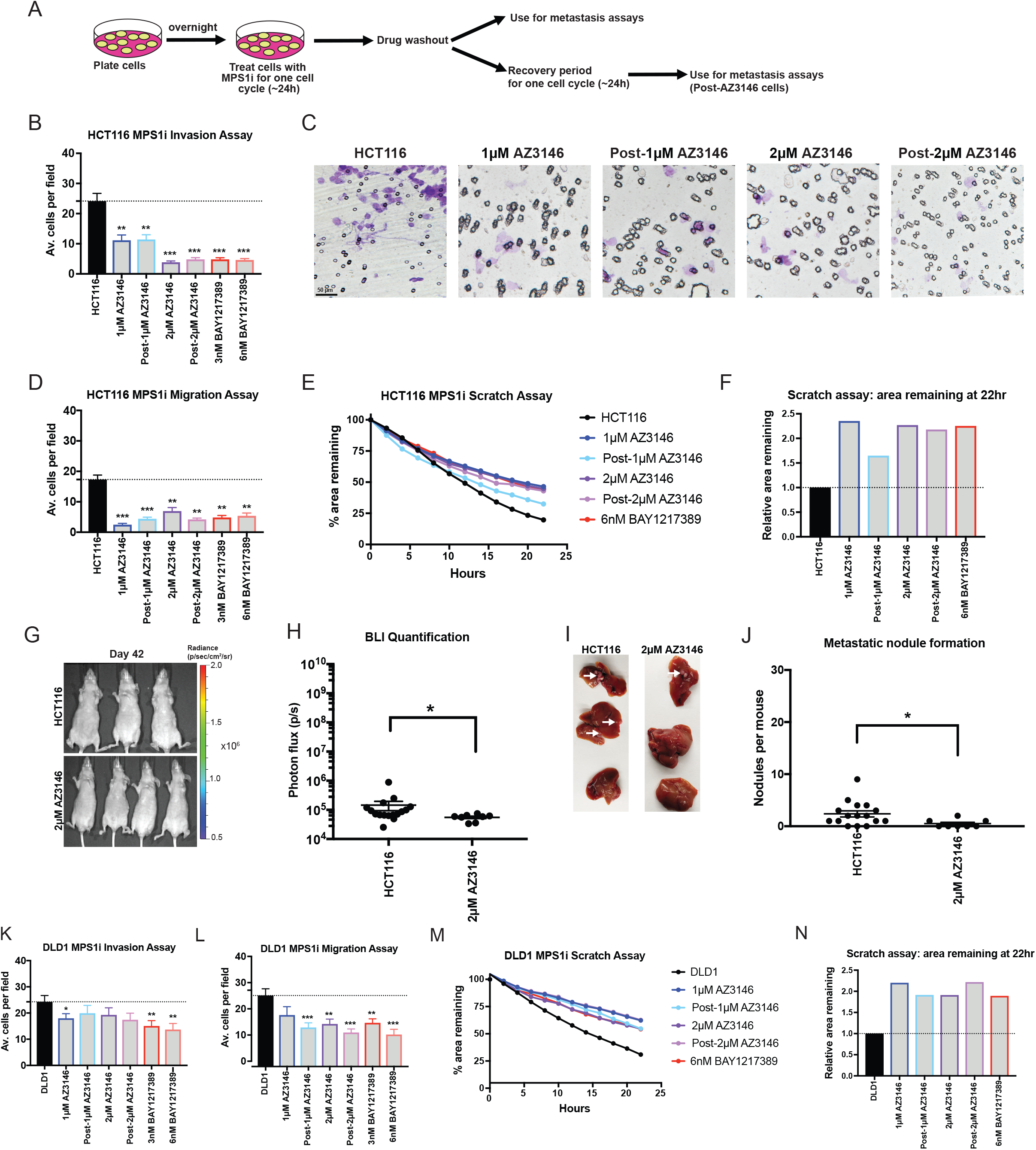
Chromosomal instability is insufficient to drive metastasis. (A) Schematic diagram of MPS1 inhibitor treatment and drug washout prior to metastasis assays. (B, K) Quantification of the average number of cells per field that were able to cross the membrane in the invasion assay in either HCT116 or DLD1. Averages represent three independent trials in which 15-20 fields were counted. (C) Representative images of invasion in the indicated Mps1i-treated cell lines. (D, L) Quantification of the average number of cells per field that were able to cross the membrane in the migration assay in either HCT116 or DLD1. (E, M) Quantification of the percent area remaining in the scratch assay after Mps1 inhibitor treatment. (F, N) The ratio of the area remaining at 22 hours after the scratch is plotted relative to the untreated parental cell line. (G) Luciferase imaging of mice injected with HCT116 or AZ3146 treated cells 42 days post-injection. (H) Bioluminescence imaging quantification of the luciferase signal in mice injected with HCT116 cells and HCT116 cells treated with 2 µM AZ3146. (I) Images of metastatic nodule formation from mice euthanized 42 days post-injection. White arrows indicate metastases. (J) Quantification of the number of nodules per mouse (HCT116 n = 16, AZ3146 n = 8). The animals injected with HCT116 cells are the same batch used in figure 1N. Bars represent mean ± SEM. Unpaired t-test * p<0.05; ** p<0.005 *** p<0.0005. Scale bar, 50 µm

Each trisomic cell line was derived via single-cell cloning, and HCT116 exhibits a very high mutation rate (Glaab and Tindall 1997). While our data indicated that gaining chromosome 5 induces a profound alteration in cell identity, it remained possible that these results were an artefact of a mutation or some other alteration acquired during the MMCT process. Notably, our observation that a single Ts5 clone (“clone 11”) retained epithelial gene expression raised the possibility that our results were due to a secondary alteration that occurred during cell line derivation. We hypothesized that, if gaining Chr. 5 caused epithelial gene silencing, then eliminating that trisomy should restore epithelial gene expression. Alternately, if this phenotype was a byproduct of a mutation acquired during cloning, then the loss of epithelial cell identity should be independent of the presence of the extra chromosome. To differentiate between these possibilities, we used two approaches to generate derivatives of HCT116 Ts5 that had lost the extra chromosome (Figure 2B). First, we grew these cells as a subcutaneous xenograft in a nude mouse, a process that we previously demonstrated selected for cells that had lost the extra chromosome (Sheltzer et al. 2017). Secondly, we cultured HCT116 Ts5 cells for several weeks to allow them to missegregate the trisomy, and then we performed a second round of single-cell cloning. Using SMASH sequencing, we confirmed that the post-xenograft cell line had lost the extra chromosome, and we identified three “evolved” clones that had lost most or all of the extra chromosome (Figure S1D). We found that all four Ts5-loss cell lines had restored EpCAM, E-cadherin, and Claudin 7 expression (Figure 2C), and this correlated with a decrease in Matrigel invasion (Figure 2D). This indicates that the prior silencing of these epithelial genes was likely a consequence of the initial aneuploidy, while the aberrant expression profile in Ts5 clone 11 is most likely due to a secondary alteration or other clonal artefact.

In addition to the EMT, cancer cell invasion is also associated with the up-regulation of matrix metalloproteinases (MMPs), a set of enzymes capable of degrading extracellular matrix components (Gialeli et al. 2011). Using qPCR, we found that HCT116 Ts5 clones up-regulated several classes of MMPs, including gelatinases (MMP2), stromelysin-like proteinases (MMP11), transmembrane MMPs (MMP14 and MMP24), and GPI-type proteinases (MMP17)(Figure S4A). However, MMPs were not uniformly up-regulated by Ts5, as we observed no consistent changes in MMP15, MMP21, or MMP25 expression. Non-invasive HCT116 Ts3 cells did not up-regulate any of the MMPs that we tested (Figure S4A).

Finally, we sought to discover whether the loss of epithelial gene expression and the up-regulation of MMPs caused the increased invasive behavior in the Ts5 cells. First, we assessed invasiveness in Ts5 clone 11 (which maintained EpCAM and E-cadherin expression), and we verified that significantly fewer cells from this clone were able to invade through a basement membrane compared to other Ts5 clones that silenced epithelial genes (Figure 2E). Next, we expressed either EpCAM or E-cadherin cDNA under a constitutive promoter in two different Ts5 clones. Western blotting revealed that these constructs led to EpCAM and E-cadherin expression levels comparable to those found in the near-diploid parental HCT116 line (Figure 2F-G). We then tested the effects of restoring EpCAM or E-cadherin expression on invasive behavior, and we found that the expression of either protein was sufficient to decrease Matrigel invasion to a level indistinguishable from the wild-type control (Figure 2H-I). Treating Ts5 cells with an MMP inhibitor also decreased Matrigel invasion, though not to wild-type levels (Figure S4B). In total, these results demonstrate that gaining an extra copy of chromosome 5 increases invasiveness in colon cancer cells by multiple mechanisms, and reestablishing epithelial gene expression is sufficient to block invasive behavior.

### Triggering chromosomal instability with Mps1 inhibitors suppresses invasive behavior

Our findings indicated that specific aneuploidies can directly affect metastasis, suggesting that the frequent appearance of aneuploidy in metastatic tumors is not simply a by-product of CIN. We therefore set out to investigate the relationship between CIN, whole-chromosome aneuploidy, and invasive behavior. In order to generate CIN, we used two inhibitors of the Mps1 kinase, AZ3146 and BAY1217389, at two different concentrations. We performed live-cell imaging in HCT116 cells that expressed H2B-GFP and we verified that treatment with either drug induced a variety of mitotic errors, including anaphase bridges, lagging chromosomes, and micronuclei, in a dose-dependent manner (Figure S5). Furthermore, after exposure to 1μM AZ3146, single-cell sequencing of HCT116 revealed that 92% of cells exhibited whole-chromosome aneuploidy, compared to only 8% of cells in an untreated population (Figure S6A-C). Thus, Mps1 inhibitors allow us to trigger transient periods of chromosomal instability, which generated populations of cells with different aneuploid karyotypes.

Next, we tested the effects of these Mps1 inhibitors on invasion and motility in HCT116 and DLD1 cells. To generate CIN, we cultured cells in low concentrations of AZ3146 or BAY1217389 for 24 hours, then washed out the drug and performed the assays described above (Figure 3A). Additionally, we performed a set of experiments in which we allowed AZ3146-treated cells to recover for 24 hours prior to commencing the assays. Surprisingly, we found that exposure to these drugs significantly suppressed invasive behavior in nearly every condition tested (Figure 3). For instance, treatment with either 2μM AZ3146 or 6nM BAY1217389 decreased Matrigel invasion in HCT116 cells by more than 80% (Figure 3B-C). These drugs also decreased pore migration and slowed the closure of a monolayer scratch (Figure 3D-F and J-M). Finally, HCT116 cells treated with 2μM AZ3146 formed significantly fewer metastatic nodules compared to the parental line when injected into the spleens of nude mice (Figure 3G-I). These results suggest that Mps1 inhibitors are capable of suppressing invasive behavior.

To confirm that these results were not specific for colon cancer cells, we performed additional sets of Matrigel invasion assays following Mps1 inhibitor treatment in the near-diploid Cal51 breast cancer cell line, the highly-aneuploid A375 melanoma cell line, and in RPE1-hTert cells. As we observed with HCT116 and DLD1 cells, blocking Mps1 significantly reduced Matrigel invasion, particularly at the higher concentration of either AZ3146 or BAY1217389 (Figure S7). Thus, in a variety of different cancer types and genetic backgrounds, exposure to a drug that increases chromosomal instability suppresses rather than enhances invasive behavior.

Many small molecules can exhibit promiscuous activity against other kinases, leading us to consider the possibility that an off-target effect of these drugs suppressed invasion (Giuliano et al. 2018; Klaeger et al. 2017). To test whether this phenotype represented an on-target effect of Mps1 inhibition, we used CRISPR-mediated homology-directed repair (HDR) to introduce two mutations into the MPS1 gene (C604W and S611G; Figure S8A, Table S1A) that have been reported to block the binding of AZ3146 to Mps1 (Gurden et al. 2015). Live-cell imaging verified that HCT116 cells harboring these mutations displayed minimal CIN when cultured in 2 μM AZ3146 (Figure S8B-C). Correspondingly, while 2 μM AZ3146 resulted in a significant decrease in Matrigel invasion in wild-type HCT116, treatment of cells harboring either Mps1^C604W^ or Mps1^S611G^ with AZ3146 had no effect on invasion (Figure S8D-E). These results demonstrate that AZ3146 suppresses invasion specifically through its inhibition of the SAC kinase Mps1.

We next considered the possibility that our experimental approach (in which cells were examined after Mps1i-washout) blocked the pro-invasive effects of CIN, perhaps by restoring normal chromosome segregation fidelity. As aneuploidy is sufficient to increase CIN (Sheltzer et al. 2011; Passerini et al. 2016), and as micronuclei that result from prior missegregation events are liable segregate incorrectly during subsequent mitoses (Soto et al. 2018), we anticipated that cells would still display instability following drug washout. Indeed, live cell imaging 24 hours after Mps1i-washout demonstrated that cells continued to exhibit a significant increase in mitotic errors compared to untreated control cells (Figure S9A). To further verify that Mps1 inhibitors decreased cellular invasion, we repeated the Matrigel invasion assays in the presence of low doses of AZ3146 or BAY1217389, without drug washout. Under these conditions, drug treatment caused a significant decrease in Matrigel invasion, verifying that increased invasiveness is not an obligate consequence of ongoing CIN (Figure S9B).

As MPS1 is an essential gene (Maciejowski et al. 2010), we next sought to test whether cellular toxicity caused by transient exposure to Mps1 inhibitors could explain the decrease in invasion that we observed. First, we assessed PARP cleavage and caspase-3 cleavage (markers of apoptosis) and beta-galactosidase staining (a marker of senescence) in cells treated with Mps1 inhibitors. 3nM BAY1217389, 6nM BAY1217389, and 1 μm AZ3146 treatment had no effect on apoptosis, while only the highest concentration of AZ3146 (2 μm) caused a slight increase in the appearance of cleaved PARP and cleaved caspase-3 (Figure S10A). Similarly, fewer than 1% of Mps1i-treated cells stained positive for senescence-associated beta-galactosidase (Figure S10B-C). Thus, under multiple conditions, Mps1 inhibitors suppress invasion without significantly increasing apoptosis or senescence. As an alternate approach to investigate this question, we exposed cells to AZ3146 treatment for 24 hours and then cultured them in drug-free media for either 7 or 14 days. We reasoned that these conditions would allow cells to recover from any acute toxicity caused by the Mps1 inhibitor. In contrast, we hypothesized that aneuploidy and cytoplasmic DNA that arise from Mps1-induced CIN may persist (though the least-fit aneuploid cells could be out-competed by other cells over time). Indeed, treatment with an Mps1 inhibitor caused an increase in the number of cells that harbored micronuclei, and this increase remained apparent up to 14 days after the drug had been washed out (Figure S10D-E; and see below). We found that HCT116 cells displayed lower levels of Matrigel invasion, even 7 or 14 days after exposure to AZ3146 (Figure S10F). Under these same conditions, cells treated with a low dose of either Mps1 inhibitor proliferated at an indistinguishable rate compared to wild-type cells (Figure S10G). In total, these experiments indicate that CIN is capable of suppressing cellular invasion, and that these results are unlikely to reflect an off-target effect or acute toxicity caused by Mps1 inhibitors.

### cGAS/STING activity is insufficient to trigger an EMT or explain the invasive behavior of HCT116 Ts5 cells

Chromosomal instability has been reported to promote invasive behavior by activating non-canonical NFκB signaling via the cytosolic DNA sensor cGAS and its adaptor STING (Bakhoum et al. 2018). As we found that Mps1 inhibitors are capable of suppressing cellular invasion, we sought to test whether they similarly affected these pathways. First, we confirmed that both AZ3146 and BAY1217389 lead to a significant, six to ten-fold increase in micronuclei in both HCT116 and DLD1 cells 24 hours after drug washout (Figure 4A-B). While the expression of STING was normally low in these cells, treatment with AZ3146 or BAY1217389 caused an increase in the appearance of cells that were strongly STING-positive (Figure 4C-D). Mps1 inhibitor treatment also caused an increase of five to eleven-fold in the translocation of the NFκB transcription factor RelB into the nucleus (Figure 4E-F). We also confirmed via qPCR that multiple NFκB targets were up-regulated (Figure 4G). Thus, these results demonstrate that the cytosolic DNA sensing pathway is functional in these cells, and that Mps1 inhibition is sufficient to activate a RelB-related transcriptional response. However, in a variety of different cancer cell lines, neither Mps1 inhibitor was sufficient to drive cellular invasion (Figure 3) or to alter the expression of the canonical EMT proteins EpCAM, E-cadherin, N-cadherin, or Vimentin (Figure S11). We conclude that an EMT is not an obligate consequence of chromosomal instability, and instead the previously-reported association between CIN, EMT, and metastasis may be limited to certain cancer types or specific ways of generating CIN.

**Figure 4.**
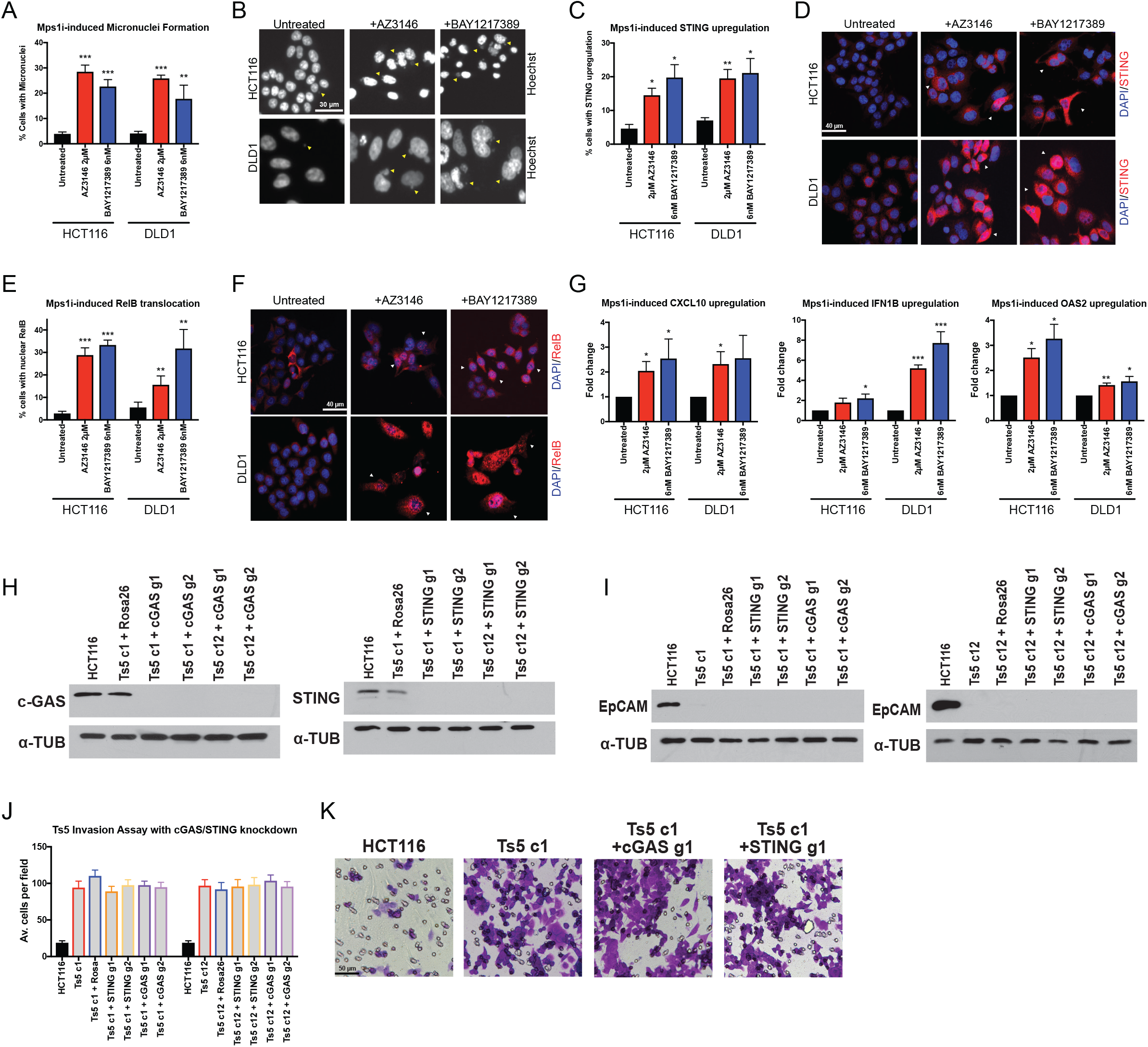
Mps1 inhibition activates cGAS/STING signaling. (A) Quantification of the percent of cells with micronuclei after Mps1 inhibitor treatment in HCT116 and DLD1. (B) Representative images of micronuclei after Mps1 inhibitor treatment. Yellow arrows indicate certain visible micronuclei. (C) Quantification of the percent of cells with STING upregulation after Mps1 inhibitor treatment. (D) Representative images of cells with STING upregulation after Mps1 inhibitors treatment. White arrows indicate certain cells with STING upregulation. (E) Quantification of the percent of cells with nuclear RelB after Mps1 inhibitor treatment. (F) Representative images of cells after Mps1 inhibitor treatment. White arrows indicate certain cells with nuclear RelB. (G) qPCR quantification of several NFκB target genes. (H) Western blot demonstrating the CRISPRi-induced knockdown of cGAS and STING in two HCT116 Ts5 clones. (I) EpCAM expression after knockdown of cGAS and STING with CRISPRi in HCT116 Ts5 clones. (J) Quantification of the average number of cells per field that were able to cross the membrane in the invasion assay after cGAS or STING knockdown. (K) Representative images of the invasion assay in the indicated cell lines. Bars represent mean ± SEM. Unpaired t-test * p<0.05; ** p<0.005 *** p<0.0005. Scale bar, 30 and 50 µm

Despite these results, cGAS/STING signaling undoubtedly has a profound effect on multiple cancer-related phenotypes. Additionally, STING is encoded on chromosome 5, and multiple inflammatory genes are upregulated in trisomic cell lines (Dürrbaum et al. 2014; Viganó et al. 2018), leading us to investigate whether cGAS/STING contributed to the metastatic behavior of the HCT116 Ts5 cells. To accomplish this, we stably expressed a dCAS9-KRAB CRISPRi vector in two independent Ts5 clones, which allowed us to trigger the specific down-regulation of a gene of interest (Horlbeck et al. 2016). We then transduced each line with two guide RNAs (gRNAs) that targeted either cGAS or STING, and we verified via western blot that these guides strongly suppressed cGAS and STING expression, respectively (Figure 4H). However, we found that silencing cGAS or STING had no effect on either EpCAM expression or invasive behavior in Ts5 (Figure 4I-K). Thus, whole-chromosome aneuploidy is capable of driving invasion in a cGAS/STING-independent manner.

We hypothesized that the over-expression of a gene or genes on chromosome 5 led to the silencing of epithelial genes and the increased invasive behavior in HCT116 Ts5. We selected 24 candidate genes encoded on chromosome 5 that had known roles in development, signal transduction, or gene regulation, and we decreased their expression with CRISPRi in two independent Ts5 clones. However, none of these knockdowns was sufficient to restore EpCAM expression (Figure S12, Table S1B-S2). The metastatic behavior of HCT116 Ts5 may therefore result from the coordinated over-expression of several Chr5 genes, or from a gene not included among this panel.

### Different chromosomal aneuploidies are associated with distinct clinical outcomes

Our results indicate that whole-chromosome aneuploidies have the capability to either promote or inhibit invasive behavior. Additionally, our previous research has demonstrated that certain aneuploidies can directly suppress transformation (Sheltzer et al. 2017). Yet, clinically, aneuploidy is widely reported to correlate with cancer progression and aggressive disease (Merkel and McGuire 1990). How can these disparate observations be reconciled?

To address this, we performed Cox proportional hazards survival modeling on 10,686 patients with 27 different types of cancer from The Cancer Genome Atlas (TCGA; other abbreviations are defined in Table S3A). In this approach, one or more clinical variables are regressed against patient outcome, allowing us to identify features that correlate with prognosis. Here, we report the Z scores from these models, which capture both the directionality and significance of a particular clinical association. If a particular chromosome gain is significantly associated with patient death, then a Z score >1.96 corresponds to a P value < 0.05 for that relationship. In contrast, a Z score less than −1.96 indicates that a chromosome gain event is significantly associated with patient survival (or that a chromosome loss event is associated with death). 10,133 of these patients’ tumors have been analyzed by SNP-chip, allowing their tumors’ bulk karyotype to be determined (Taylor et al. 2018). Tumors in the TCGA lack comprehensive annotation of metastases, so we instead used either “overall survival” or “progression-free survival” as clinical endpoints for this analysis (Table S3A, and see the Materials and Methods).

First, we regressed each tumor’s total aneuploidy burden against patient outcome. We determined that highly-aneuploid cancers were associated with a significantly worse prognosis than tumors with low aneuploidy in nine of 27 different patient cohorts (Figure 5A-B). In the remaining 18 cancer types, no significant association was detected. We did not find any cohorts in which high aneuploidy portended a favorable outcome (Figure 5B). Some cancer types in which aneuploidy has previously been reported to be a prognostic biomarker were not found to exhibit a significant correlation in this analysis (e.g., lung cancer; Choma et al. 2001); this may be because of differences in the technology used to measure aneuploidy, differences between patient populations, or because of the limited number of patients included in some cohorts. The acquisition of aneuploidy was linked to the loss of p53 function, as p53-mutant tumors tended to have higher levels of aneuploidy than p53-wild-type tumors (Figure S13A). However, the overlap between p53 mutations and highly-aneuploid tumors was unable to fully account for the prognostic significance of aneuploidy: in multivariate Cox regression models that included both p53 status and aneuploidy, high aneuploidy burden was still significantly correlated with poor survival in six different cohorts (Figure S13B-C).

**Figure 5.**
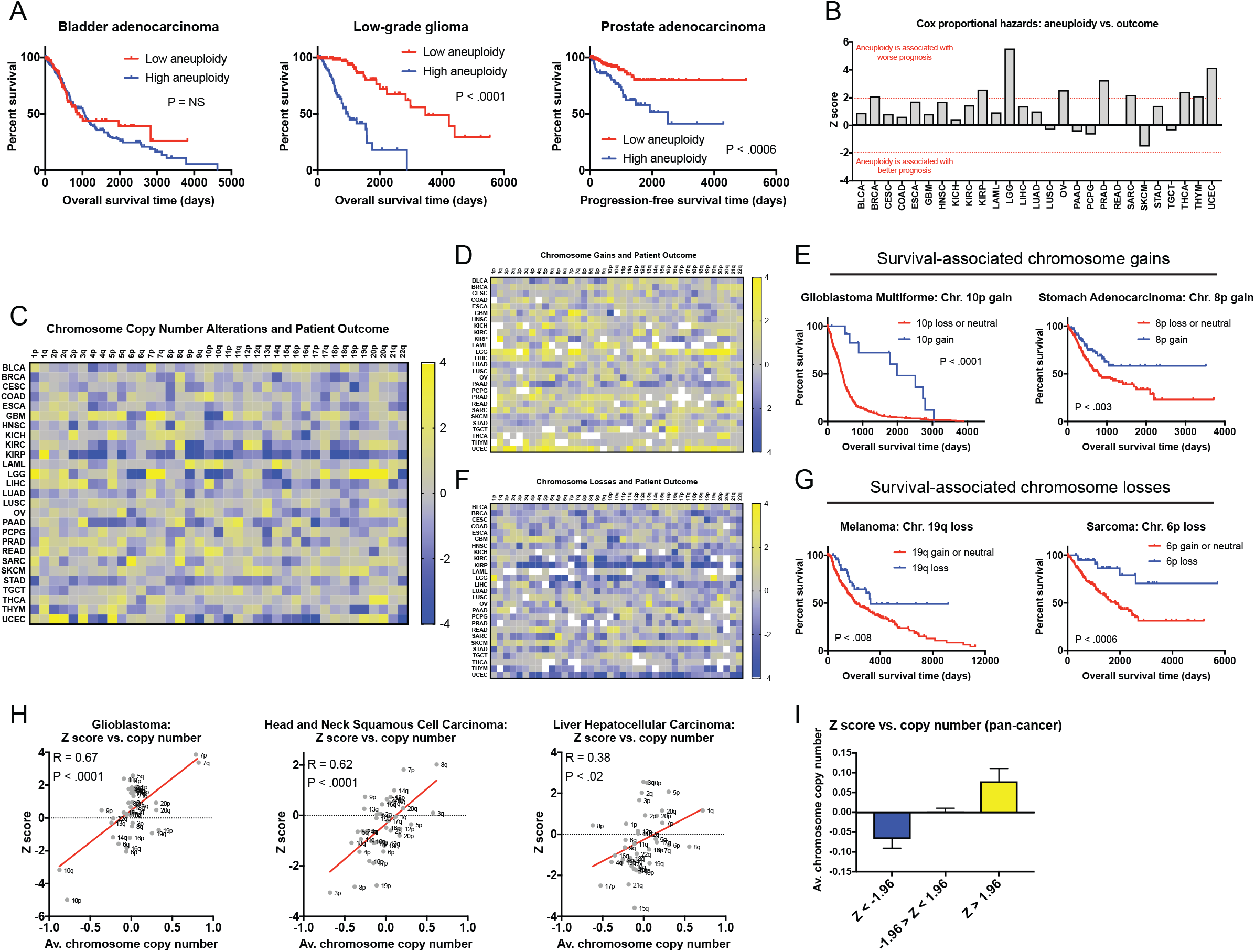
A pan-cancer analysis of chromosome copy number changes associated with patient survival. (A) The total number of aneuploid chromosomes in each tumor was calculated according to (Taylor et al. 2018). In each cancer type, patients were subdivided into two groups: low aneuploidy (≤ 20^th^ percentile aneuploidy score) and high aneuploidy (≥ 80^th^ percentile aneuploidy score). Kaplan-Meier curves are shown for three cancer types. (B) Z scores from Cox-proportional hazards analysis based on the aneuploidy scores, as described above, are displayed. Note that Z > 1.96 indicates a significant association between higher aneuploidy and death, while Z < −1.96 indicates a significant association between higher aneuploidy and survival. (C) A heatmap comparing chromosome arm copy number vs. patient outcome across 10,133 patients with 27 different types of cancer. The color-bar indicates the Z score from the Cox analysis. For visualization purposes, Z scores were capped at 4 and −4. (D) A heatmap comparing dichotomized chromosome arm copy number vs. patient outcome across 10,133 patients with 27 different types of cancer. In this analysis, survival was compared between patients in which a given arm was gained and patients in which a given arm was either lost or was copy-neutral. The color-bar indicates the Z score from the Cox analysis. For visualization purposes, Z scores were capped at 4 and −4. (E) Representative Kaplan-Meier curves of two instances in which chromosome gains are associated with improved patient prognosis. (F) A heatmap comparing dichotomized chromosome arm copy number vs. patient outcome across 10,133 patients with 27 different types of cancer. In this analysis, survival was compared between patients in which a given arm was lost and patients in which a given arm was either gained or was copy-neutral. The color-bar indicates the Z score from the Cox analysis. For visualization purposes, Z scores were capped at 4 and −4. (G) Representative Kaplan-Meier curves of two instances in which chromosome losses are associated with improved patient prognosis. (H) Scatter plots comparing the average chromosome copy number vs. the Z score obtained from Cox analysis for three different cancer types. (I) A bar graph displaying the average chromosome copy number across all 27 cancer types, binned based on the Z score from the Cox model.

We next set out to determine how specific aneuploidies influenced patient prognosis. We constructed Cox models to interrogate the relationship between the alteration status of each chromosome arm (“loss”, “neutral”, or “gain”) and patient outcome across the 27 cohorts. This analysis revealed 160 arm-length aneuploidies that were associated with survival time (Figure 5C and Table S3B). In every cancer type, at least one arm-length alteration was associated with outcome, indicating that specific aneuploidies can be prognostic factors even in cancer types in which bulk aneuploidy is uninformative. However, no aneuploidy was associated with outcome in more than six of 27 cancer types. Thus, there is no single “pro-metastatic” karyotype, and the effects of specific aneuploidies on cancer cell dissemination likely depend on a tumor’s genetic and epigenetic background.

As our *in cellulo* and *in vivo* work had primarily focused on colon cancer, we next examined the clinical correlates of aneuploidy in this patient cohort in particular. We found that total aneuploidy tended to increase with colon cancer stage: stage IV tumors harbored a median of 16 arm-length alterations, nearly twice as many as the median stage I or II tumor (Figure S14A). Interestingly, colon cancer aneuploidy was not significantly associated with a worse response to therapy (Figure S14B). However, tumors that expressed high levels of an EMT-related gene signature displayed more aneuploidy than tumors that lacked this signature, suggesting a potential *in vivo* link between aneuploidy and cell state transitions (Figure S14C). Colon tumors commonly displayed a number of recurrent aneuploidies, including the gain of Chr20q, Chr7, and Chr13, and the loss of Chr18 and Chr17 (Figure S14D-E). We did not detect a significant correlation between the gain of chromosome 5 and patient outcome; as RPE1 Ts5 cells also did not display a significant increase in invasiveness, this is consistent with our hypothesis that aneuploidies interact with a cell’s genetic or epigenetic background to drive metastasis (Figure S14F). In fact, no whole-chromosome events were individually associated with colon cancer outcome, and only three arm-length alterations were (8p, 10p, and 17q; Figure S14F-G). Thus, as we observed in our cell line models, a majority of individual aneuploidies are not strong drivers of tumor metastasis.

Across all 27 cancer cohorts, we observed a positive relationship between chromosome copy number and survival (Z > 1.96) for 66 arm-length aneuploidies, while we found a negative relation (Z < −1.96) for 94 aneuploidies. These results could be consistent with two different models: a positive Z score could indicate that tumors that harbor an amplification of that chromosome have a worse prognosis than tumors that lack that amplification, or it could indicate that tumors that have lost a copy of that chromosome have a better prognosis than tumors that are copy-neutral. Similarly, negative Z scores could indicate that chromosome losses accelerate patient death or that chromosome gains protect against it. To differentiate between these models, we re-analyzed the TCGA survival and karyotype data, binning “loss” and “neutral” calls together, or binning “gain” and “neutral” calls together (Figure 5 D-G and Table S3C-D). This analysis revealed that in ∼87% of cases, the aneuploidy event (either gain or loss) was associated with worse patient outcome. However, strikingly, in ∼13% of cases, the gain or loss of a chromosome arm correlated with improved patient survival. For instance, glioblastomas that are copy-neutral or have lost Chr10p have a median survival time of 422 days, while glioblastomas that harbor a gain of Chr10p have a median survival time of 1987 days (Figure 5E). Thus, while most prognostic aneuploidies correlate with poor outcomes, aneuploidy of many different individual chromosomes can be associated with decreased tumor aggressiveness.

As an alternate proxy for tumor aggressiveness, we repeated the above analysis, using a patient’s “progression-free interval” (PFI) as an endpoint. This endpoint is defined as the time until a patient exhibits loco-regional or systemic recurrence, a second malignancy, or death from any cause (Liu et al. 2018). Similar to the results that we obtained when using “overall survival” as an endpoint, we uncovered nine cancer types in which high aneuploidy was associated with poor PFI, and we did not observe any cases in which high aneuploidy correlated with prolonged PFI (Figure S15A). We identified 182 significant correlations between specific aneuploidies and PFI, which included at least one significant relationship in every cancer lineage (Figure S15B and Table S3E). 88% of individual aneuploidies were associated with worse PFI, while 12% of aneuploidies were associated with prolonged PFI (Figure S15C-D and Table S3F-G).

If aneuploidy is capable of either promoting or suppressing aggressive behavior, then why is bulk tumor aneuploidy a common hallmark of poor prognosis? To address this, we compared the arm-level Z scores obtained from our survival analysis with the overall frequency of each aneuploidy event. We found that, in general, chromosome arms that were associated with poor prognosis when amplified were significantly more likely to be gained than lost in primary tumors. Similarly, arms whose deletions correlated with dismal survival were more commonly lost than gained (Figure 5H-I). For instance, in head and neck squamous cell carcinoma, 8q amplifications are associated with poor prognosis, and this chromosome arm was gained in 54% of patients and lost in <1% of patients. In contrast, 3p deletions are a biomarker of poor prognosis, and this chromosome arm was lost in 61% of patients and gained in <1% (Figure 5H). This suggests that aneuploidies capable of suppressing aggressive behavior are selected against during tumor development, leading to an enrichment of metastasis-promoting aneuploidies, even though they represent only a fraction of all possible copy number changes.

## Discussion

Stoichiometric imbalances in endogenous proteins caused by aneuploidy interfere with multiple cellular functions (Sheltzer and Amon 2011). It is therefore conceivable that cell motility, matrix degradation, and the other processes necessary for invasive behavior might also be perturbed by aneuploidy-induced dosage imbalances. Thus, while aneuploidy is a hallmark of cancer, aneuploidy in itself might potently suppress certain functions otherwise needed for tumorigenicity and metastasis. To investigate the relationship between aneuploidy and metastasis, we have generated and analyzed a series of modified cell lines that differ from each other by a single chromosome. Using this system, we discovered that single-chromosome gains can have multifaceted effects on several proxies for metastatic ability. Across 13 different trisomies, 12 inhibited or had a minor effect in assays designed to test invasiveness and cell motility.

HCT116 colon cancer cells harboring an extra copy of chromosome 5 behaved differently from every other trisomy studied and displayed a clear increase in metastatic capacity. We traced this phenotype to an aneuploidy-induced partial epithelial-mesenchymal transition, in which the specific amplification of chromosome 5 causes the silencing of epithelial cell adhesion genes. This effect is independent of cGAS/STING signaling, and we posit that the over-expression of one or more genes on chromosome 5 cooperate to induce this phenotypic switch. Aneuploidy-induced cell state transitions may be a novel cause of the strong correlation between aneuploidy and death from cancer. Additionally, recent evidence has underscored the importance of phenotypic plasticity in the metastatic cascade and has suggested that an EMT is neither a necessary nor an irreversible step in cancer cell dissemination (Chen et al. 2019; Padmanaban et al. 2019; Zheng et al. 2015; Fischer et al. 2015; Liu et al. 2019). Instead, cancer cells can lose certain epithelial characteristics to promote migration out of the primary tumor site and into circulation, but they can re-establish them (through a mesenchymal-epithelial transition) in order to colonize a distant location (Dongre and Weinberg 2019). Aneuploidy, too, can be reversible: gaining and losing chromosomes through CIN could contribute to this plasticity and allow cancer cells to switch between different phenotypic states.

CIN has previously been reported to directly influence metastasis by triggering a cGAS/STING-dependent EMT and increase in cell motility (Bakhoum et al, 2018). By generating CIN with Mps1 inhibitors, we have verified the link between chromosomal instability and cGAS/STING activation, but we find that these inhibitors suppress rather than promote metastatic behavior. These results demonstrate that increased invasiveness is not an obligate consequence of either CIN or cGAS/STING signaling and may instead depend on the mechanism by which CIN arises. As CIN causes the acquisition of aneuploidy, we believe that, as with the single-chromosome trisomic cell lines that we have characterized, CIN-induced proteome deregulation may interfere with the processes necessary to achieve a metastasis-competent state. Low levels of CIN, or CIN that allows the development of rare, metastasis-promoting aneuploidies, may be necessary to observe the conditions under which CIN enhances metastasis. Additionally, as several Mps1 inhibitors have entered clinical trials in human patients (Xie et al. 2017), our results further suggest that one potential benefit of Mps1 inhibition as a therapeutic strategy may be an overall decrease in metastatic dissemination.

Interestingly, while Ts5 enhanced metastasis in HCT116 cells, this same chromosome mildly suppressed metastatic behavior when added to retinal-pigment epithelial cells. We speculate that the gain of chromosome 5 has a specific effect on HCT116 cells based on their genetic and epigenetic background, and our pan-cancer analysis of aneuploidy-associated patient mortality supports this hypothesis. Many aneuploidies exhibit distinct clinical associations in different cancer types: for instance, Chr10p losses are associated with poor survival in gliomas and kidney cancers, but Chr10p amplifications are associated with poor survival in leukemia and hepatocellular carcinoma (Table S3B). We further observed that, among amplified chromosomes, Chr8q gains showed the strongest correlation with patient death, but this association was only present in six of 27 cancer types (Table S3B). We find no evidence to support the hypothesis that specific aneuploidies are universal metastasis promoters (Duesberg et al. 2006). Instead, the consequences of each aneuploidy are closely tied to the original tumor type. Yet, chromosome copy number changes can provide raw fodder for tumor evolution. While a subset of aneuploidies do in fact correlate with decreased aggressiveness in patient samples, these aneuploidies are rarely observed. The copy number changes that instead drive malignant behavior are selected through evolution, and thus “deadly” aneuploidies appear to be more common than they actually are. These results reconcile the apparent contradiction between the behavior of artificially-constructed aneuploid cells, which almost always display a pronounced growth defect, and the aggressive behavior of aneuploid human tumors (Stingele et al. 2012; Sheltzer et al. 2017).

## Methods

### Sample sizes and statistical methodology

For the Matrigel invasion assays (Figure 1B, Figure 2D, Figure 2E, Figure 2H, Figure 3B, Figure 3K, Figure 4J, Figure S2A, Figure S2F, Figure S4B, Figure S8D, Figure S9C, and Figure S10F) we analyzed 15-20 independent fields of view, collected from 2 to 4 independent biological replicates of the experiment. For the migration assays (Figure 1E, Figure 3D, Figure 3L, Figure S2C, and Figure S2H) we analyzed 15-20 independent fields of view, collected from 2 to 4 independent biological replicates of the experiment. For the scratch assays (Figure 1G, Figure 3E, Figure 3M, Figure S2D, and Figure S2I) we analyzed 8-10 fields along the scratch, collected from 2-4 independent biological replicates of the experiment. For the spleen-liver metastasis assays (Figure 1N, Figure 3J), 16 NU/J mice were injected with HCT116 luciferase-expressing cells, 10 mice were injected with HCT116 Ts5 c1 luciferase-expressing cells, 8 mice were injected with HCT116 Ts3 luciferase-expressing cells, and 8 mice were injected with HCT116 luciferase-expressing cells treated with 2 µM AZ3146. For Mps1i-induced micronuclei formation (Figure 4A and Figure S10D), we analyzed a minimum of 500 cells per trial collected from 3 independent biological replicates of the experiment. For the STING and Rel-B immunofluorescence assays (Figure 4C and Figure 4E), we analyzed a minimum of 500 cells per trial collected from 3 independent biological replicates of the experiment. For the qPCR assays (Figure 4G, Figure S4A, and Figure S12B), we performed 3 technical replicates collected from 3 independent biological replicates of the experiment. For live cell mitotic imaging (Figure S5A, Figure S8B, and Figure S9A), we analyzed a minimum of 100 cells from 1-2 independent biological replicates of the experiment.

For the invasion, migration, scratch, immunofluorescence, and live-cell imaging experiments, the samples were analyzed while blinded to cell identity. No outliers were excluded from analysis. Bar graphs display sample means +/-SEM. Unpaired t-tests were used to measure statistical significance.

### Cell lines and tissue culture conditions

The identity of each human cell line was verified by STR profiling (University of Arizona Genetics Core, Tucson, AZ). The karyotype of every aneuploid cell line used in this manuscript was verified with the SMASH technique (Wang et al. 2016) or had been previously analyzed with a similar low-pass whole-genome sequencing method (Sheltzer et al. 2017). The names of the clones and their karyotypes are summarized in Table S4. HCT116, DLD1, RPE1, Cal51, SW480, and A375 were grown in DMEM supplemented with 10% FBS, 2 mM glutamine, and 100 U/mL penicillin and streptomycin. T47D cells were grown in RPMI supplemented with 10% FBS, 6.94 μg/ml insulin (Thermo Fisher, Waltham, MA; BN00226), 2 mM glutamine, and 100 U/ml penicillin and streptomycin. A549 cells were grown in Ham F12 supplemented with 10% FBS, 100 U/mL penicillin and streptomycin and 2 mM glutamine. All cell lines were grown in a humidified environment at 37°C and 5% CO_2_.

### Karyotype analysis with SMASH

Cells were trypsinized and resuspended in PBS, centrifuged at 1000 rpm for 5 minutes, and then the pellets were collected. Total cellular genomic DNA was isolated using the Qiagen QIAamp kit (Cat. No. 51036). SMASH karyotyping was performed as described (Wang et al. 2016). In brief, total cell genomic DNA was enzymatically fragmented to a mean size of ∼40bp and joined to create chimeric fragments of DNA suitable for creating NGS libraries (300-700bp). The fragment size selection was done with Agencourt AMPure XP beads (Beckman Coulter, Cat. No. A63881). Illumina-compatible NEBNext Multiplex Dual Index Primer Pairs and adapters (New England Biolabs, Cat. No. E6440S) were ligated to the selected chimeric DNA fragments. These barcoded DNA fragments were then sequenced using an Illumina MiSeq. Bin boundaries were determined empirically by apportioning valid SMASH mappings from 56 CHD trios sequenced at high depth to a fixed number of bins such that the minimum bin count is maximized and the remaining bins are populated as evenly as possible (Andrews et al. 2016; Andrews 2017). Approximately 2% of the resulting bins showed significant variation in count across the CHD population and were thus excluded from further analysis and display using both automated means and human review. The generated reads were demultiplexed and mapped using custom scripts, and plots were generated with G-Graph (Andrews 2017).

### Single-cell sequencing

This protocol was adapted from (Baslan et al. 2012). In brief, HCT116 cells were plated into 6 well cell culture plates and 24hr later the cells were treated with 1μM AZ3146. The next day, the media was changed. 24hr later, the cells were trypsinized and single-cell sorted into 96 well PCR plates (ThermoFisher Scientific, Cat. No. AB0731) containing cell lysis buffer (0.1% SDS, 2% Triton) and incubated at 65°C for 1 hour. Genomic DNA was enzymatically digested with NlaIII (NEB, Cat. No. R0125). Both ends of fragments were tagged by using oligonucleotides that contain cell-barcode, universal primer and several random nucleotides (varietal tags) through ligation and extension reactions. Cell barcodes allow us to multiplex samples, and the tags allow unique counting of initial DNA fragments. After amplification by universal primer, fragment size selection was performed with Agencourt AMPure XP beads (Beckman Coulter, A63881) to obtain fragments appropriate for sequencing. Barcoded sequencing adapters were ligated to the DNA fragments which allows us to further multiplex samples for sequencing. Next generation sequencing was done on an Illumina MiSeq. Reads were demultiplexed using customized scripts and karyotype plots were generated with G-Graph (Andrews 2017).

### Invasion and migration assays

For transwell migration assays, cells were plated in the upper chamber of the transwell insert (Corning Cat. No. 3464, 24-well insert, pore size: 8 µm). For DLD1 and HCT116 cells, 3 × 10^4^ cells were seeded, while for RPE1 cells 1× 10^4^ cells were seeded. For transwell invasion assays, cells were plated in the top chamber with Matrigel-coated membranes (Corning Cat. No. 354480, 24-well insert, pore size: 8 µm). For DLD1, 1 × 10^5^ cells were seeded, for HCT116, 2 × 10^5^ cells were seeded, and for RPE1, 1.5 × 10^4^ cells were seeded. Cells in the upper chamber were plated in serum-free DMEM, while media with 10% FBS was added to the lower chamber. The plates were incubated for 24 hours at 37°C, and then the cells on the upper surface were removed using cotton swabs. The membranes were then fixed in methanol and stained with crystal violet dye. The membranes were cut out and mounted on to slides. They were imaged at 40x (15-20 images) and counted to obtain the average number of cells per field that migrated or invaded. Two to three chambers were used per cell line and/or condition. For experiments using Mps1 inhibitors, the cells were either treated for 24 hours with the indicated drug, followed by 24 hours of drug-free washout prior to seeding, or the cells were treated with the indicated drug for 24 hours and then seeded immediately.

### Scratch assays

For scratch assays, 1-1.5 × 10^6^ cells were seeded on a 6-well dish and 24 hours later (at around 90% confluency) the monolayer was scratched with a 200 µL pipette tip. The cell culture plate was then placed in an inverted Zeiss Observer for live-cell imaging (5% CO_2_, 37°C). Phase contrast images were taken every 30 mins for 22 h at 8-10 positions along the scratch and 2-3 trials were repeated for each cell line and condition. Data was analyzed with ImageJ to determine the scratch area remaining using the Wound Healing Tool Macro (Collins 2007). For experiments with Mps1 inhibitors, the cells were either treated for 24 hours with the indicated drug followed by 24 hours of drug-free recovery prior to seeding, or the cells were treated with the indicated drug for 24 hours immediately after seeding. A minimum of 2 trials per cell line and treatment were performed.

### Western blot analysis

Whole cell lysates were harvested and resuspended in RIPA buffer [25 mM Tris, pH 7.4, 150 mM NaCl, 1% Triton X 100, 0.5% sodium deoxycholate, 0.1% sodium dodecyl sulfate, protease inhibitor cocktail (Sigma, Cat. No. 4693159001), and phosphatase inhibitor cocktail (Sigma, Cat. No. 4906845001)]. Quantification of protein concentration was done using the RC DC Protein Assay (Bio-Rad, Hercules, CA; Cat. No. 500–0119). Equal amounts of lysate were denatured and loaded onto a 10% SDS-PAGE gel. The Trans-Blot Turbo Transfer System (Bio-Rad) and polyvinylidene difluoride membranes were used for protein transfer. Antibody blocking was done with 5% milk in TBST (19 mM Tris base, NaCl 137 mM, KCl 2.7 mM and 0.1% Tween-20) for 1 hour at room temperature except for E-cadherin, cGAS, N-cadherin and Vimentin which used 5% BSA in TBST. The following antibodies and dilutions were used: E-cadherin (Cell Signal, Danvers, MA; Cat. No. 3195) at a dilution of 1:1000 (5% BSA), N-cadherin (Cell Signal; Cat. No. 13116) at a dilution of 1:1000 (5% BSA), Vimentin (Cell Signal; Cat. No. 5741) at a dilution of 1:1000 (5% BSA), EpCAM (Abcam, Cambridge, MA; Cat. No. ab124825) at a dilution of 1:1000 (5% milk), Claudin-7 (Abcam; Cat. No. ab27487) at a dilution of 1:1000 (5% milk), Fibronectin (Abcam; Cat. No. ab32419) at a dilution of 1:1000 (5% milk), cGAS (Sigma-Aldrich; Cat. No. HPA031700) at a dilution of 1:1000 (5% BSA), STING (Cell Signal; Cat. No. 13647) at a dilution of 1:2000 (5% milk), cleaved PARP (Cell Signal; Cat. No. 5625) at a dilution of 1:1000 (5% milk), and cleaved Caspase 3 (Cell Signal; Cat. No. 9661) at a dilution of 1:1000 (5% milk). Blots were incubated with the primary antibody overnight at 4°C. Anti-alpha tubulin (Sigma-Aldrich; Cat. No. T6199) at a dilution of 1:20,000 was used as a loading control. Membranes were washed at RT 3 times (10 mins each) before they were incubated in secondary antibodies for an hour at RT. HRP goat anti-mouse (Bio-Rad; Cat. No. 1706516) at 1:50,000 was used for tubulin blots while HRP goat anti-rabbit (Abcam; Cat. No. ab6721) at 1:30,000 was used for all other primary antibodies. Membranes were washed 3 times again (15 min each) and developed using ProtoGlow ECl (National Diagnostics; Cat. No. CL-300) and autoradiographic film (Lab Scientific; XAR ALF 2025).

### Overexpression of E-Cadherin and EpCAM

The following plasmids were obtained from Applied Biological Materials (Richmond, BC, Canada): E-cadherin (Cat. No. LV704934), EpCAM (Cat. No. LV149412) and an empty control plasmid (Cat. No. LV591). The plasmids were transfected into HCT116 Trisomy 5 cells using Lipofectamine 3000 (ThermoFisher, Cat. No. L3000001). The cells were then selected with puromycin for 2 weeks and then bulk-sorted with FACS for RFP-expressing cells. Invasion assays and western blot analysis was performed on cells that were expanded following FACS purification.

### Senescence staining

To test for senescence, cells were stained for beta-galactosidase using the Cell Signaling Technology kit (#9860) as per the manufacturer’s protocol. As a positive control, cells were treated with etoposide (20 μM) for 24 hours followed by 4 days of recovery. A minimum of 500 cells were counted per condition.

### CRISPRi plasmid construction and virus generation

Guide RNAs for CRISPRi experiments were chosen from (Horlbeck et al. 2016). Guides were cloned into the LRG2.1 mCherry vector (Addgene; Cat. No. 108099) using a BsmBI digestion as previously described (Giuliano et al. 2019). Plasmids were transformed in Stbl3 *E. coli* (Thermo Fisher; Cat. No. C737303) and sequenced to confirm the presence of the correct gRNA sequence. A dCas9-KRAB construct (Addgene; Cat. No. 85969) was used to knock down target gene expression. Guide RNA sequences are listed in Table S1A.

Lentivirus was generated using calcium phosphate transfection as previously described (Chang et al. 2013). Supernatant was harvested at 2 to 3 intervals 48 to 72 hr post-transfection by filtering through a 0.45 μm syringe, and then frozen at −80° C for later use or supplied directly to cells with 4 μg/mL polybrene.

### RNA isolation and quantitative real-time PCR

Total cellular RNA was extracted and isolated using TRIzol (Life Technologies; Cat. No. 15596018) and a Qiagen RNeasy Mini Kit (Cat. No. 74106). Total RNA was converted to cDNA using SuperScript^TM^ III First-Strand System kit (ThermoFisher Scientific; Cat. No. 18080051). Quantitative PCR was performed for the target genes using SYBR Premier Ex Taq (Takara; Cat No. RR420L) and quantified using the QuantStudio 6 Flex Real-Time PCR system (Applied Biosystems). Primers are listed in Table S2.

### Immunofluorescence and Microscopy

Cells were placed on autoclaved 12mm x 12mm glass coverslips in twelve well cell culture plates. The next day, the cells were treated with the indicated drug in separate cell culture wells. 24 hours later, fresh culture media was added for a post-drug recovery period of 24 hours. Later they were fixed with either 4% paraformaldehyde for 15 min at RT (when staining for of STING) or cold (−20 ° C) methanol for 5min at RT (when staining for Rel-B). For STING, selective plasma membrane permeabilization was done using 0.02% Saponin in TBS for 2 min. For Rel-B, nuclear permeabilization was done using 0.15% of Triton X-100 in TBS for 10 min. TBS + 1% BSA + 22.52 mg/ml of glycine was used as a blocking agent for 45 minutes. TBS + 1%BSA was used as blocking agent during primary antibody staining (STING at a dilution of 1:1000; Abcam Cat. No. ab181125; RelB at a dilution of 1:500; Abcam Cat. No. ab180127) overnight at 4 ^◦^C. After washing the cells, they were incubated with goat anti-rabbit Alexa Fluor 647 (at a dilution of 1:1000; Abcam Cat. No. ab150083). The coverslips were treated with DAPI (0.1ug/ml in TBS) for 2 min. The coverslips were then mounted with Prolong Diamond Antifade Mountant (ThermoFisher Scientific; Cat No., P36965). The cells were viewed on a spinning disk confocal microscope (UltraVIEW Vox; PerkinElmer) and quantified using Volocity version 6.3.

### Live cell imaging

HCT116 cells expressing H2B-GFP were seeded and treated with Mps1 inhibitors as indicated. Live-cell imaging was performed at room temperature using spinning-disc confocal microscopy system (UltraVIEW Vox; PerkinElmer) and a charged-coupled device camera (ORCA-R2; Hamamatsu Photonics) fitted to an inverted microscope (DMI6000 B; Leica) equipped with a motorized piezoelectric stage (Applied Scientific Instrumentation). Overnight imaging was performed using a Plan Apochromat 20X 0.7 NA air objective with camera binning set to 2×2. Image acquisition and analysis was performed using Volocity version 6.3 (PerkinElmer). A minimum of 100 cells per condition were analyzed for mitotic errors. Cells were tracked form nuclear envelope breakdown to anaphase exit and were analyzed for errors such as lagging chromosomes. polar chromosomes, anaphase bridges, and multipolar mitoses.

### Quantification of micronuclei

Cells were seeded onto twelve well cell-culture plates. The next day, the cells were treated with Mps1 inhibitors as indicated. Cells were then allowed to recover in drug-free media for 24 hours. Following this period, the cells were stained with 5 ng/ml Hoescht 33342 (Invitrogen; Cat. No. H3570) for 20min at 37^◦^C. The cells were viewed under a Nikon Eclipse Ti-S microscope and quantified using NIS elements BR version 4.40. A minimum of 400 cells per condition were analyzed for micronuclei. In every field of view imaged, the total number of cells and the total number of cells with micronuclei were counted to determine the percent of cells with micronuclei.

### Knocking-in AZ3146-resistance mutations with CRISPR-mediated HDR

Mps1-targeting guides and single-stranded donor templates to introduce the AZ3146-resistance mutations were designed using Benchling (www.benchling.com). In addition to the C604W and S611G alterations, multiple silent mutations were included in the donor oligo to prevent re-cutting following template-mediated repair (Table S1A). Guide RNAs were cloned into the Lenti-Cas9-gRNA-GFP vector (Addgene # 124770) as previously described (Giuliano et al. 2019). To perform CRISPR-mediated HDR, 2μg of Mps1 gRNA plasmid was transfected along with 100 pmol of ssODN into Cas9-expressing cell lines using Lipofectamine 3000 (Thermo Fischer Scientific, Cat. No. L3000015). Successful knock-in was confirmed using the primers listed in Table S1A.

### TCGA data analysis

Survival data for TCGA patients was acquired from (Liu et al. 2018). P53 mutation data for TCGA patients was acquired from (Bailey et al. 2018). The EMT gene signature was acquired from (Gibbons and Creighton 2018). Across the 33 available TCGA cohorts, we first eliminated the six cohorts with fewer than 100 patients. Next, we chose “overall survival time” as a default endpoint, as it reflects an objective and unambiguous event. However, we noted that five of the remaining cohorts had fewer than 15 deaths; for those five cohorts we used “progression-free survival” as a clinical endpoint rather than “overall survival” (Table S3A). Additionally, several of the analyses described below were repeated using “progression-free interval” as an endpoint (Figure S15 and Table S3E-G).

Patient survival was analyzed using Cox proportional hazards modeling. The Cox model is given by the following function:

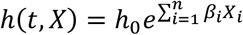

In this model, t is the survival time, h(t, X) is the hazard function, h_0_(t) is the baseline hazard, X_i_ is a prognostic variable, and β_i_ indicates the strength of the association between a variable and survival. In this model, patients have a baseline, time-dependent risk of death [h0(t)], and this risk is modified by time-independent prognostic features that either increase (β_i_>0) or decrease (β_i_<0) the patient’s risk of death. In this paper, we report Z scores, which are calculated by dividing the regression coefficient (βi) by its standard error. If a chromosome gain is significantly associated with patient death, then a Z score >1.96 corresponds to a P value < 0.05. In contrast, a Z score less than −1.96 indicates that a chromosome gain is significantly associated with patient survival (or, alternately, that a chromosome loss event is associated with death). Cox proportional hazards models and their application to TCGA data is discussed in more detail in (Smith and Sheltzer 2018).

Karyotype data for TCGA patients was acquired from (Taylor et al. 2018). Aneuploidy calls were made based on Affymetrix SNP 6.0 profiling of tumor samples. Tumor ploidy was determined using ABSOLUTE and was used to determine the baseline copy number for each patient (Carter et al. 2012). Arms or chromosomes in which more than 80% of the region was affected by a CNA were called positive for an arm or whole-chromosome alteration, while arms or chromosomes in which less than 20% of the region was affected by a CNA were considered to be negative. Aneuploidy scores were calculated by summing the total number of arm-length alterations detected in a tumor sample.

For analysis including p53, only tumors that harbored a non-synonymous mutation were considered p53-mutant (missense, non-sense, or frameshift). To assess the link between bulk aneuploidy and patient outcome, we compared survival between “low aneuploidy” tumors (≤ 20th% aneuploidy score) and “high aneuploidy” tumors (≥ 80th% aneuploidy score). Cox proportional hazards modeling was performed using Python, pandas, rpy2 and the R survival package. Code written for this analysis is available at https://github.com/joan-smith/aneuploidy-survival. This analysis also relied on packaged code from (Smith and Sheltzer 2018), now also available at https://github.com/joan-smith/biomarker-survival for ease of reuse.

### Animals and splenic injections

Nude mice (NU/J female, 6-week old; JAX stock #002019) were obtained from the Jackson Laboratory. The Cold Spring Harbor Laboratory IACUC reviewed and approved all procedures. HCT116 and trisomic cells were injected via splenic injection to generate a preclinical model of hepatic metastasis. The cell lines were transduced to express luciferase (Addgene #75020) for bioluminescence imaging.

Anaesthetized mice were swabbed with an alcoholic solution of iodine or chlorhexidine before a small incision (10-15mm) was made. The spleen was identified and carefully manipulated through the incision to sit outside the mouse on a moist gauze swab. 10^6^ cells in 100 µl PBS suspensions were injected into the organ. For the spleen, effective injection was monitored by bleaching of the organ and lack of bleeding. In some cases, splenectomy can be performed by clamping and cauterizing the splenic arteries and venous supply, thus effectively removing the spleen. The process of splenectomy carries a significant risk of bleeding, which typically occurs if the blood supply is not cauterized correctly. Successful splenic injections were verified by visual assessment for minimal bleeding. The abdominal muscle wall was then closed using an absorbable suture. The incision was closed in the top two layers using a continuous vicryl suture (Matric 1.5, Ethicon) for the peritoneum and a wound clip. Analgesia was performed with meloxicam (5-10-mg/kg PO or SQ) or Ketoprofen (5mg/kg, SC) at daily intervals for 3 days as necessary. Animals were allowed to recover from anesthesia in pre-warmed clean cages for a period of 20 minutes.

Following the injections, mice were imaged once a week for 7 weeks to monitor for tumor growth. For the imaging, mice were injected intra-peritoneally with XenoLight D-Luciferin - K+ Salt Bioluminescent Substrate (15 mg/ml, 100ul/10g; Perkin Elmer). The mice were imaged with Xenogen IVUS Spectrum (Perkin Elmer). 42 days post-injection, the mice were euthanized and metastatic nodules were visually scored.

## Acknowledgments

We thank members of the Sheltzer Lab for helpful comments on this work. Research in the Sheltzer Lab is supported by an NIH Early Independence Award (1DP5OD021385), a Breast Cancer Alliance Young Investigator Award, a Damon Runyon-Rachleff Innovation Award, a Gates Foundation Innovative Technology Solutions grant, and a CSHL-Northwell Translational Cancer Research Grant. This work was performed with assistance from the CSHL Flow Cytometry Shared Resource, which is supported by the CSHL Cancer Center Support Grant 5P30CA045508. Research in the Wigler Lab was supported by grants to M. Wigler from the Simons Foundation (SFARI 497800) and Life Sciences Founders Directed Giving-Research (award numbers 519054).

## Supplemental Figure Legends

**Figure S1.**
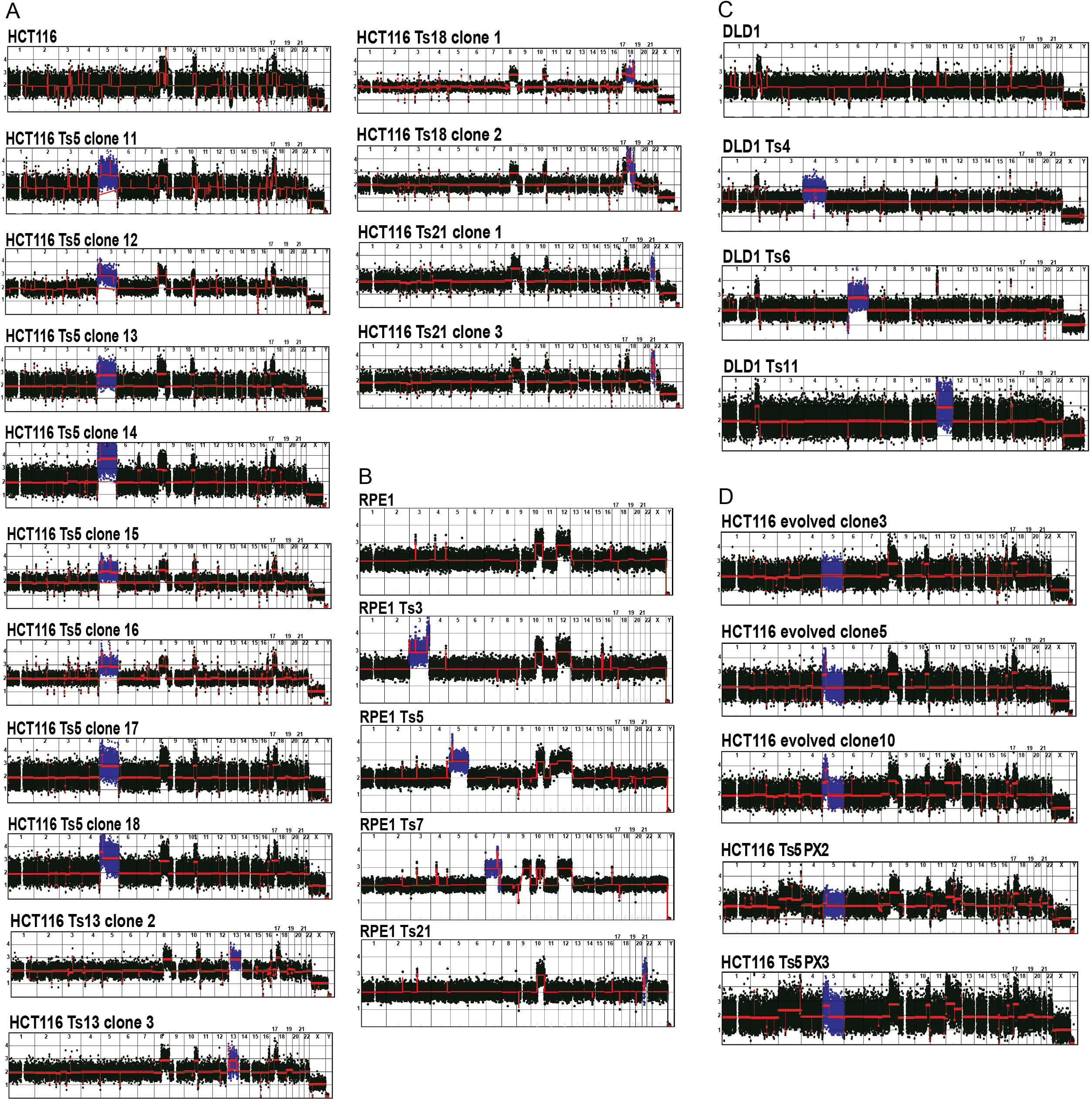
Karyotypes of the cell lines used in this study. Karyotypes were analyzed using the SMASH technique, and normalized read depths across 50kb bins are shown (Wang et al. 2016). (A) Karyotypes of HCT116 and the HCT116 trisomies. (B) Karyotypes of RPE and the RPE1 trisomies. (C) Karyotypes of DLD and the DLD1 trisomies. (D) Karyotypes obtained after reversion of HCT116 Ts5. Only evolved clone 3 and Ts5 PX2 (post-xeno) showed complete loss of both arms of chromosome 5.

**Figure S2.**
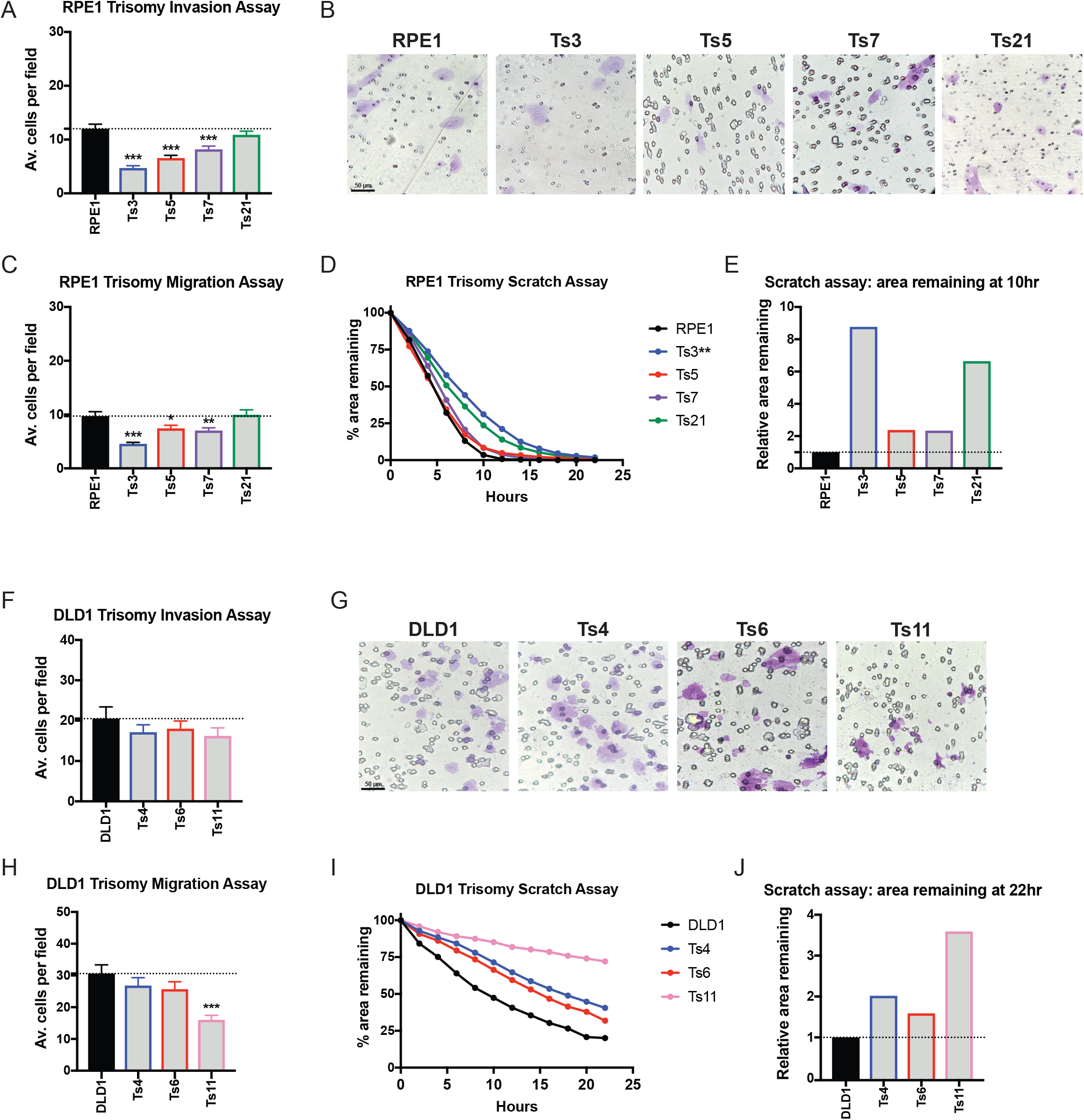
Single chromosome gains in RPE1 and DLD1 commonly suppress metastatic behavior. (A, F) Quantification of the average number of cells per field that were able to cross the membrane in the invasion assay in RPE1 and DLD1. Averages represent three independent trials in which 15-20 fields were counted. (B, G) Representative images of RPE1 and DLD1 invasion. (C, H) Quantification of the average number of cells per field that were able to cross the membrane in the migration assay. Averages represent three independent trials in which 15-20 fields were counted. (D, I) Quantification of cell motility in a scratch assay. The percent area remaining between two monolayers separated by a pipette tip-induced scratch was monitored for 22 hours. (E, J) The ratio of the area remaining at 22 hours after the scratch is plotted relative to the RPE1 or DLD1 parental cell line. A ratio less than 1 indicates faster scratch closure relative to wild-type, while a ratio greater than 1 indicates slower scratch closure. Bars represent mean ± SEM. Unpaired t-test * p<0.05; ** p<0.005 *** p<0.0005. Scale bar, 50 µm

**Figure S3.**
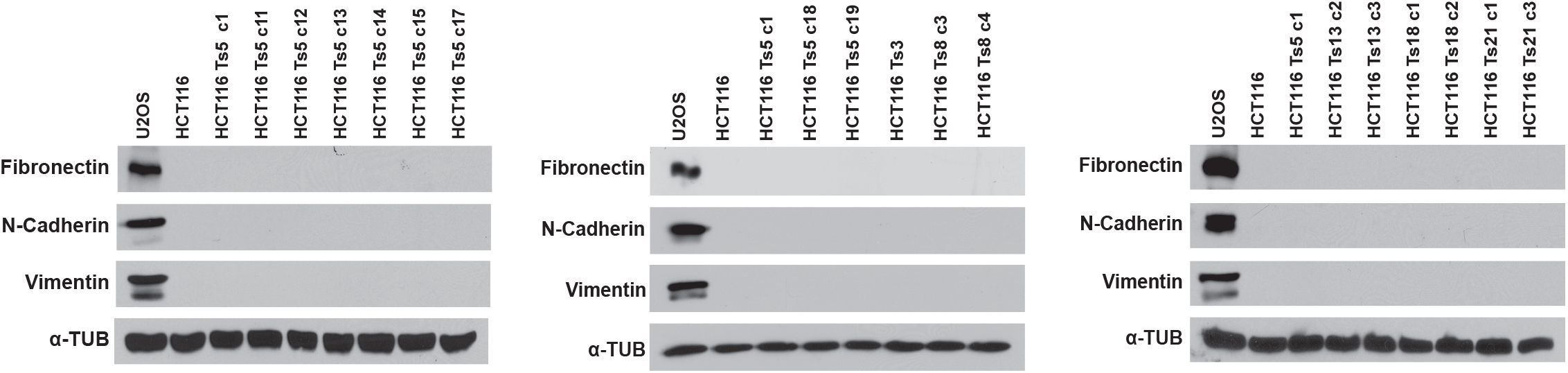
HCT116 Trisomy 5 induces a partial EMT phenotype. Western blot analysis for Fibronectin, Vimentin, and N-cadherin indicates that HCT116 trisomies do not express mesenchymal genes. The sarcoma cell line U2OS was analyzed as a positive control.

**Figure S4.**
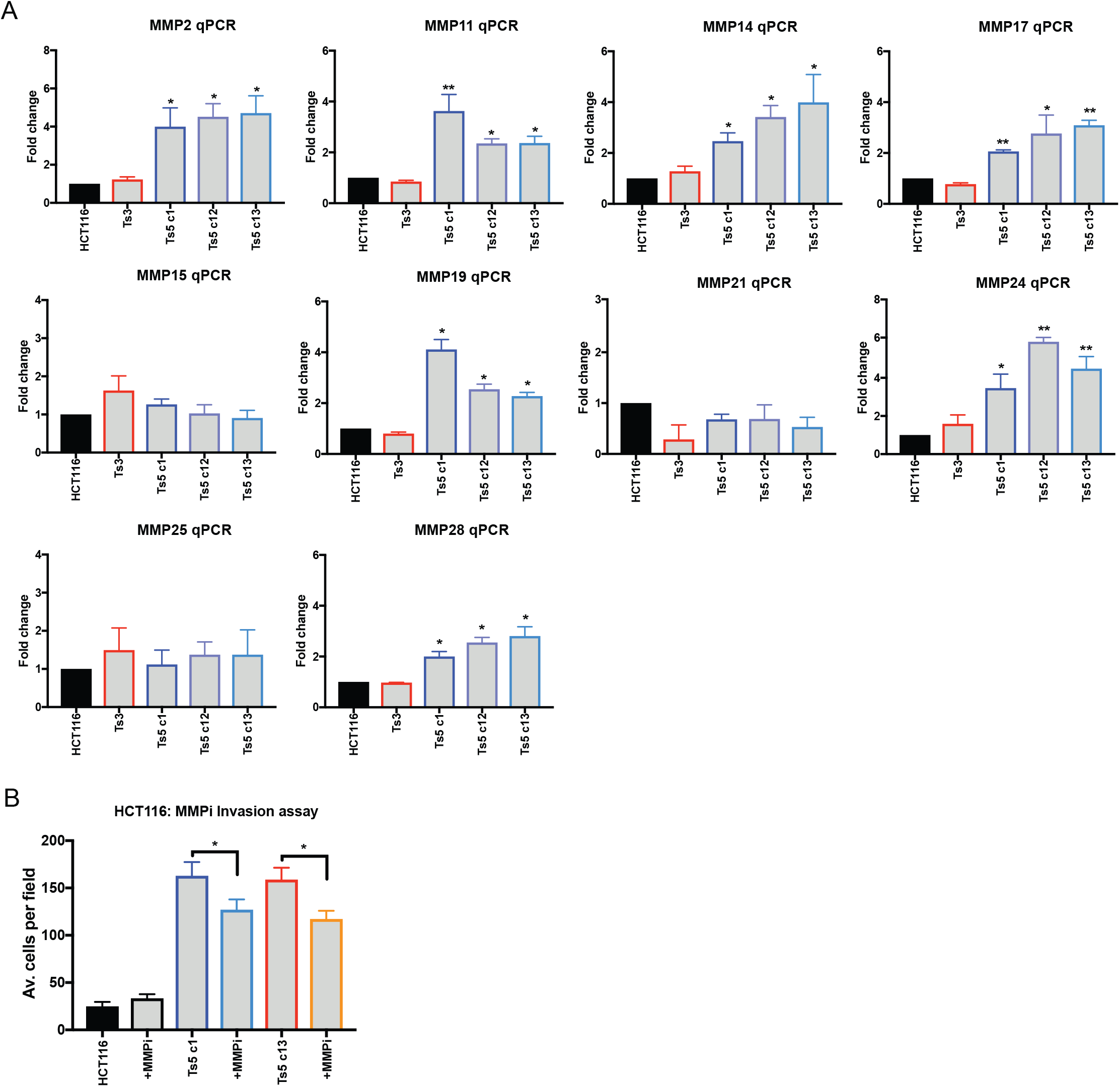
Gaining chromosome 5 causes an upregulation of MMP expression that increases cell invasiveness. (A) Quantification of several MMPs in HCT116, HCT116 Ts3, and several HCT116 Ts5 clones. (B) Treatment with 10 µM of the MMP inhibitor Batimastat decreases Matrigel invasion in HCT116 Ts5 but not in the HCT116 parental cell line. Bars represent mean ± SEM. Unpaired t-test * p<0.05; ** p<0.005 *** p<0.0005.

**Figure S5.**
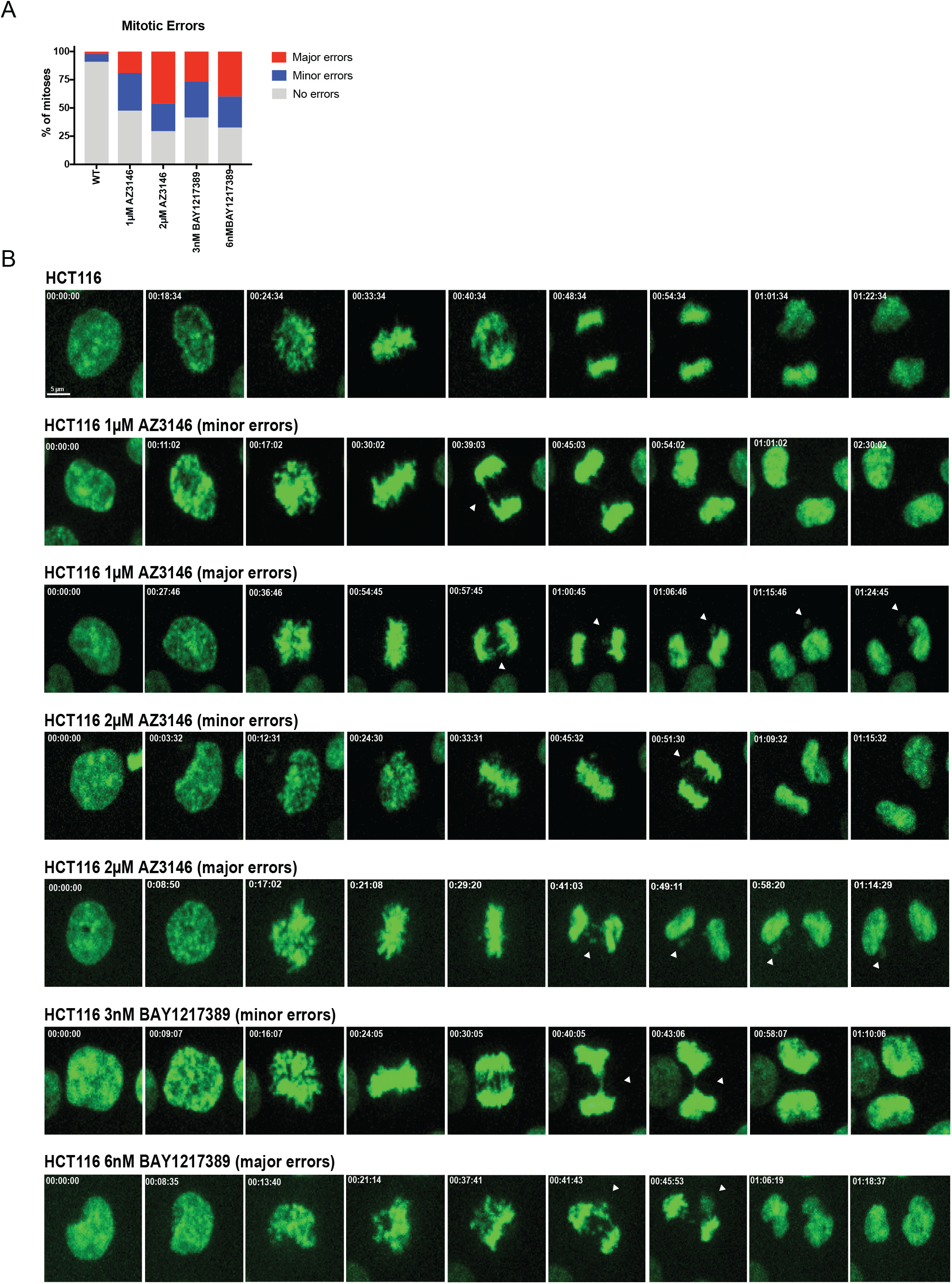
Mps1 inhibitor treatment causes random chromosome segregation errors. (A) Quantification of mitotic error frequency after Mps1 inhibitor treatment. A mitosis with “minor errors” exhibited a single lagging chromosome, anaphase bridge, or micronucleus, while a mitosis with “major errors” displayed more than one of these phenomena. (B) Representative images of chromosome segregation errors due during mitosis. White arrows indicate chromosome segregation errors. Scale bar, 5 µm.

**Figure S6.**
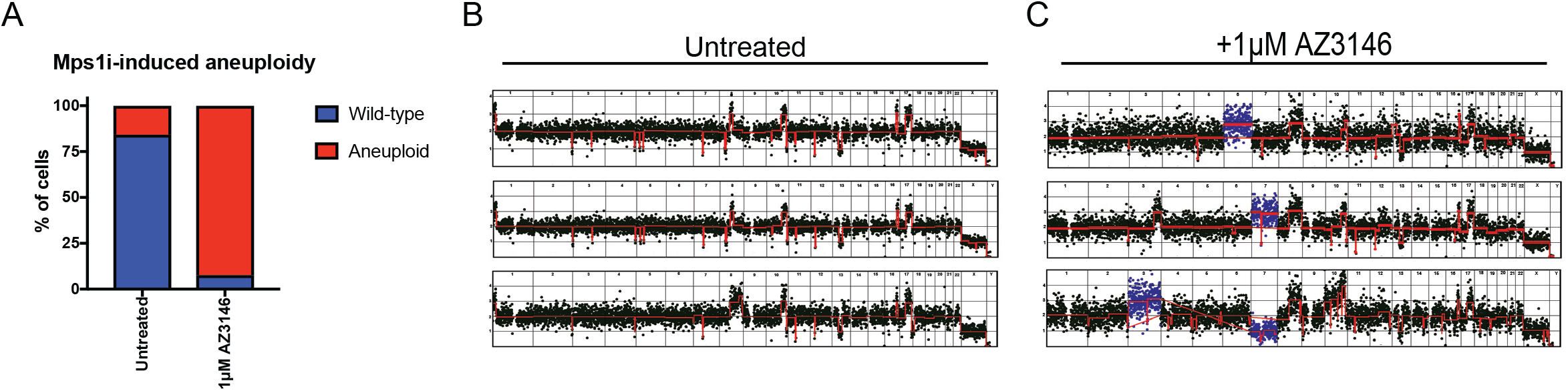
AZ3146 treatment causes whole chromosome gains and/or losses. (A) The percent of cells with whole-chromosome aneuploidies in HCT116 cells +/-1 µM AZ3146 are displayed. (B-C) Representative karyotypes of single cells +/-1 µM AZ3146.

**Figure S7.**
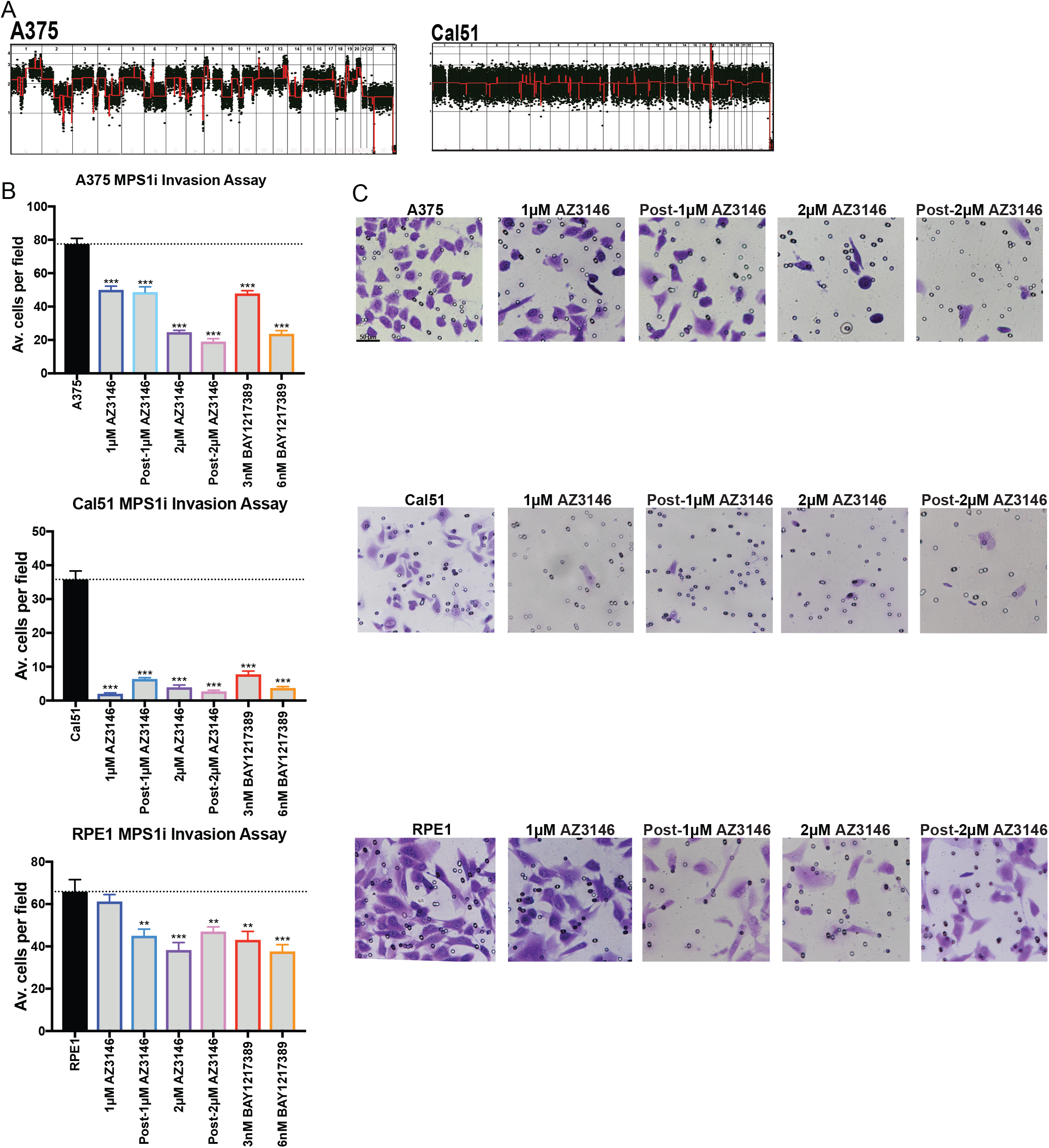
MPS1i treatment suppresses invasive behavior in multiple cell lines. (A) Karyotypes of A375 and Cal51 cell lines. (B) Quantification of the average number of cells per field that were able to cross the membrane in the invasion assay in A375, Cal51 and RPE1. Averages represent two independent trials in which 15-20 fields were counted. (C) Representative images of invasion in the indicated Mps1i-treated cell lines. Bars represent mean ± SEM. Unpaired t-test * p<0.05; ** p<0.005 *** p<0.0005. Scale bar, 50 µm

**Figure S8.**
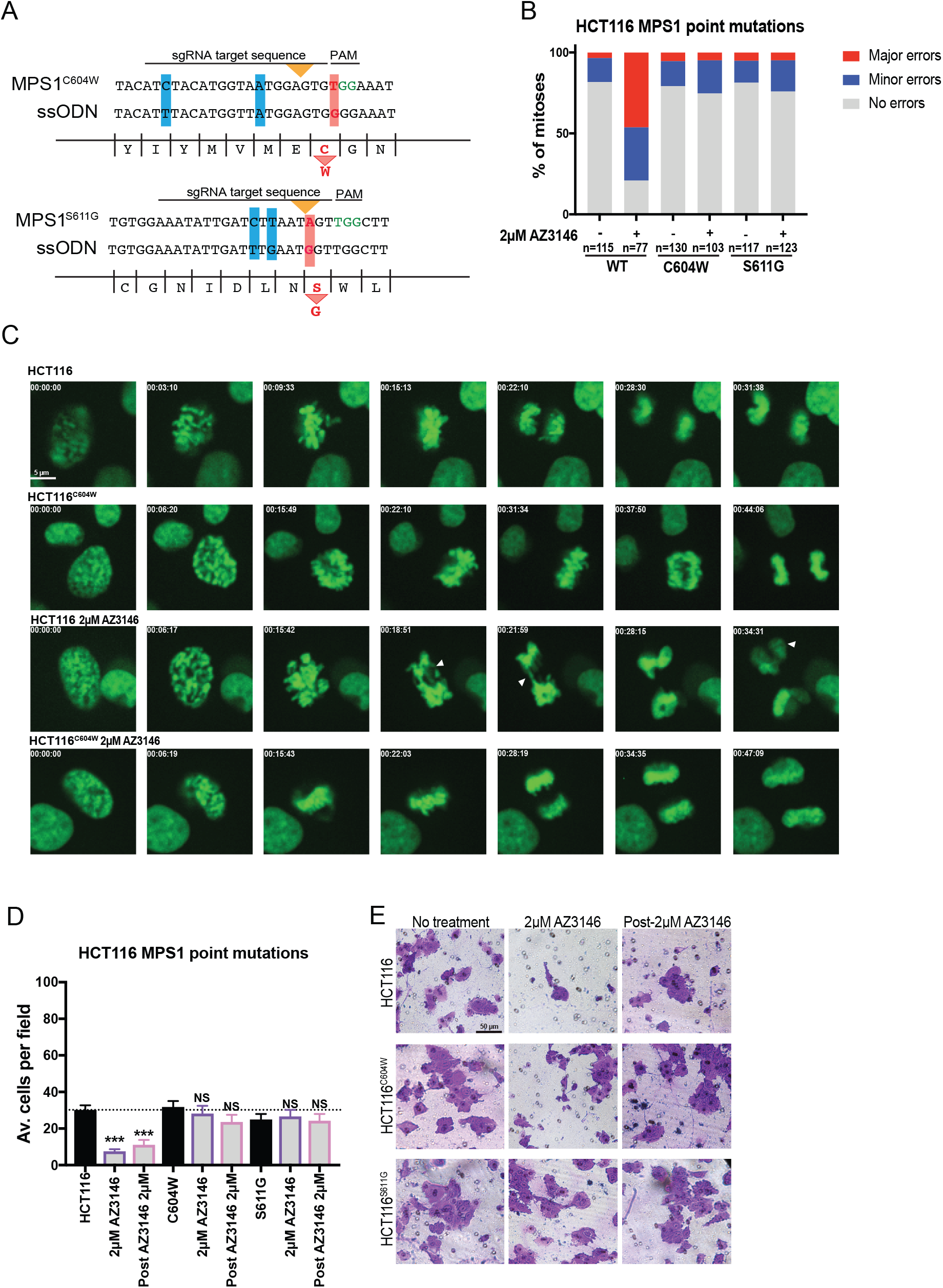
Mutations in Mps1 block the effects of AZ3146 on CIN and invasion. (A) A diagram of the CRISPR-mediated HDR strategy to introduce a C604W mutation and a S611G mutation into the endogenous MPS1 locus. Red bars highlight the missense mutations while blue bars highlight silent mutations introduced to minimize re-cutting. (B) Quantification of mitotic error frequency in the presence and absence of the Mps1 inhibitor AZ3146. A mitosis with “minor errors” exhibited a single lagging chromosome anaphase bridge, or micronucleus, while a mitosis with “major errors” displayed more than one of these phenomena. (C) Representative images of mitoses in the indicated cell lines. (D) Quantification of the average number of cells per field that were able to cross the membrane in the invasion assay in HCT116. Averages represent two independent trials in which 15-20 fields were counted. (E) Representative images of HCT116 invasion. Bars represent mean ± SEM. Unpaired t-test * p<0.05; ** p<0.005 *** p<0.0005. Scale bar, 5 and 50 µm.

**Figure S9.**
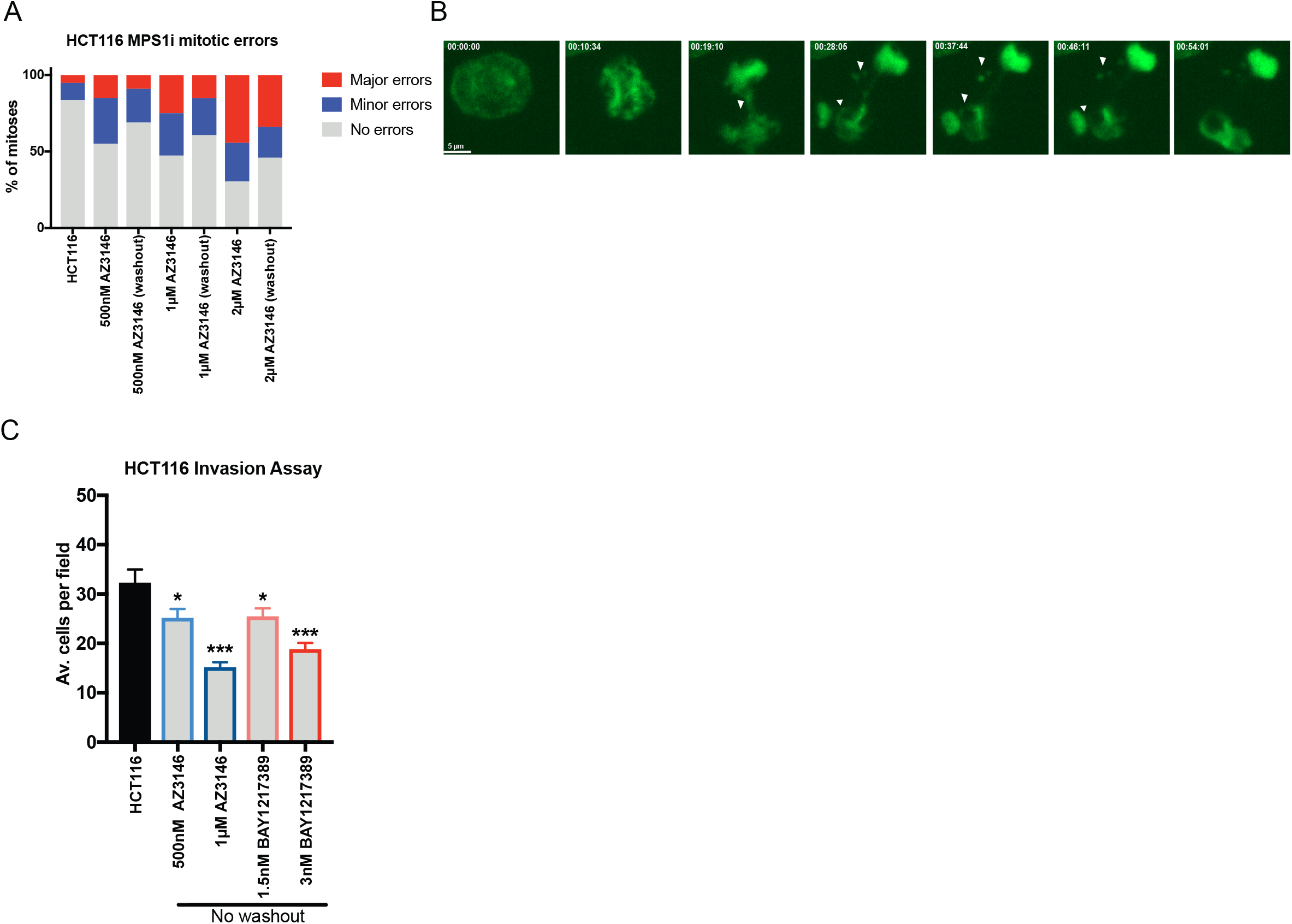
On-going CIN due to MPS1 inhibitor treatment causes random chromosome segregation errors and suppresses invasion. (A) Quantification of mitotic error frequency in the presence of the Mps1 inhibitor AZ3146 or after a 24-hour recovery period following drug-washout. A mitosis with “minor errors” exhibited a single lagging chromosome anaphase bridge, or micronucleus, while a mitosis with “major errors” displayed more than one of these phenomena. (B) Representative images of chromosome segregation errors due during a mitosis in cells that were previously treated with 1 µM AZ3146 and then allowed to recover for 24 hours in drug-free media. White arrows indicate chromosome segregation errors. (C) Quantification of the average number of cells per field that were able to cross the membrane in the invasion assay in the presence of low concentrations of Mps1 inhibitor. Note that in these experiments, the Mps1 inhibitor was included in the serum-free media at the top of the invasion chamber (Figure 1A). Averages represent two independent trials in which 15-20 fields were counted. Bars represent mean ± SEM. Unpaired t-test * p<0.05; ** p<0.005 *** p<0.0005. Scale bar, 5 µm.

**Figure S10.**
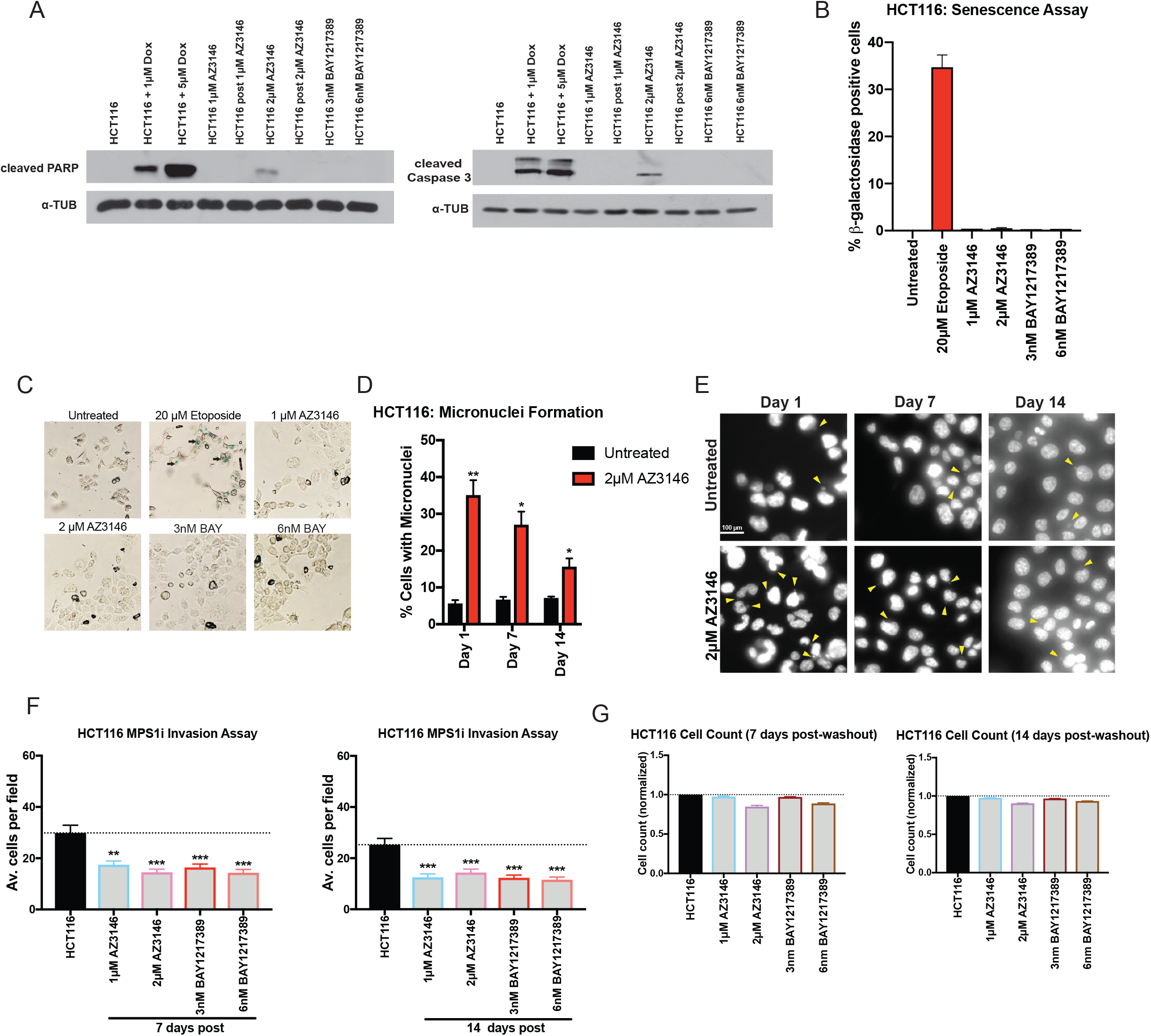
Transient exposure to Mps1 inhibitors induces minimal cellular toxicity. (A) Western blots for two markers of apoptosis, cleaved PARP and cleaved caspase-3, in Mps1i-treated cells are displayed. The DNA-damaging agent doxorubicin is used as a positive control. (B) Quantification of senescence-associated beta-galactosidase expression in Mps1i-treated cells. The DNA-damaging agent etoposide is used as a positive control. (C) Representative images of the senescence assay. Black arrows indicate blue-staining senescent cells. (D) Quantification of micronuclei formation at various time points after AZ3146 treatment and washout. (E) Representative images of HCT116 cells stained with Hoechst dye at various time points after AZ3146 treatment and washout. (F) Quantification of Matrigel invasion in HCT116 cells after treatment with AZ3146 or BAY1217389 followed by recovery in drug-free media for 7 or 14 days. (G) Quantification of cell counts in HCT116 cells after treatment with AZ3146 or BAY1217389 followed by recovery in drug-free media for 7 or 14 days. Bars represent mean ± SEM. Unpaired t-test * p<0.05; ** p<0.005 *** p<0.0005. Scale bar, 100 µm.

**Figure S11.**
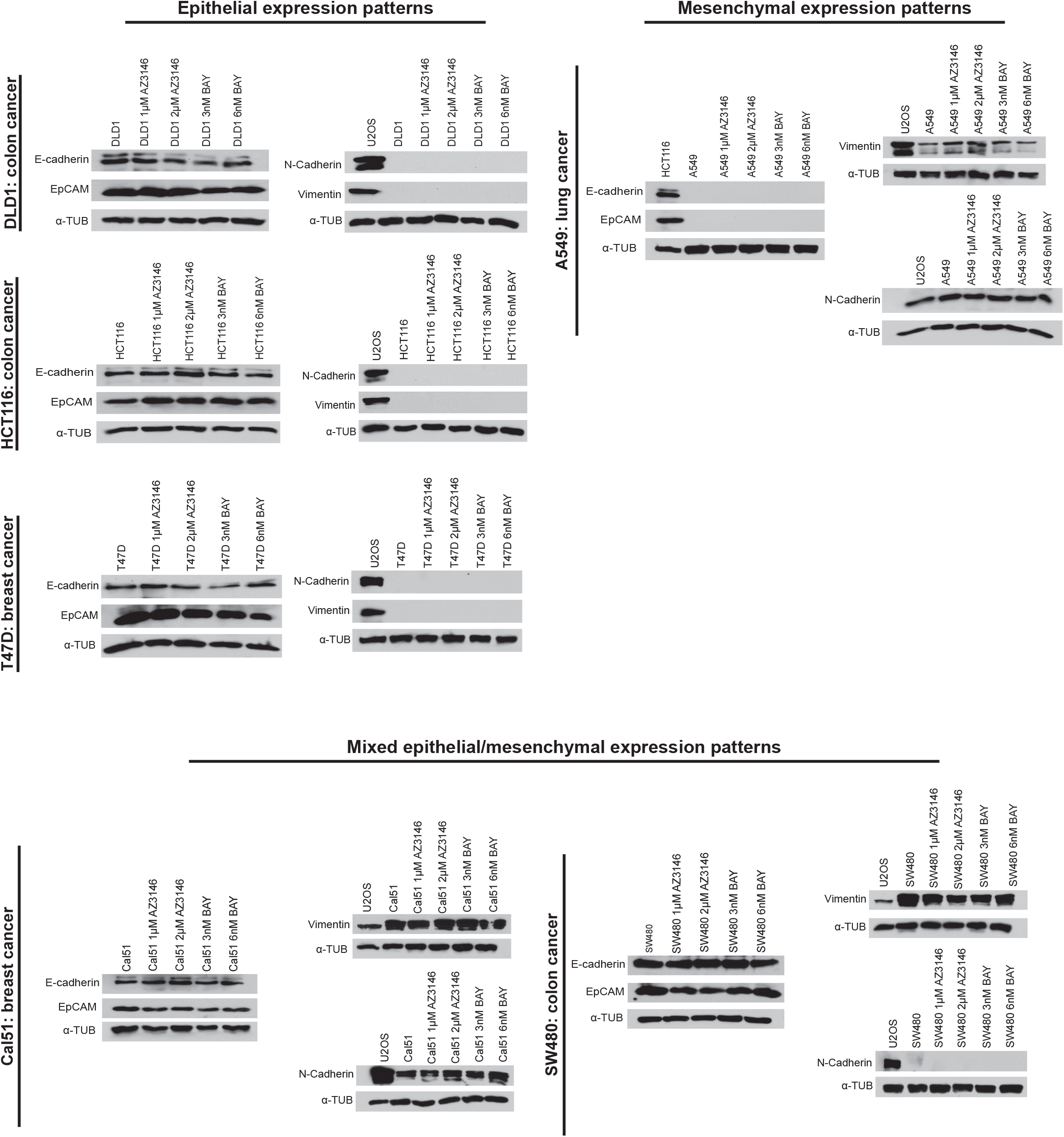
Mps1 inhibitor treatment does not affect the expression of epithelial or mesenchymal genes. Six different cancer cell lines were treated with Mps1 inhibitors and then analyzed via western blotting. The sarcoma cell line U2OS serves as a positive control for mesenchymal gene expression.

**Figure S12.**
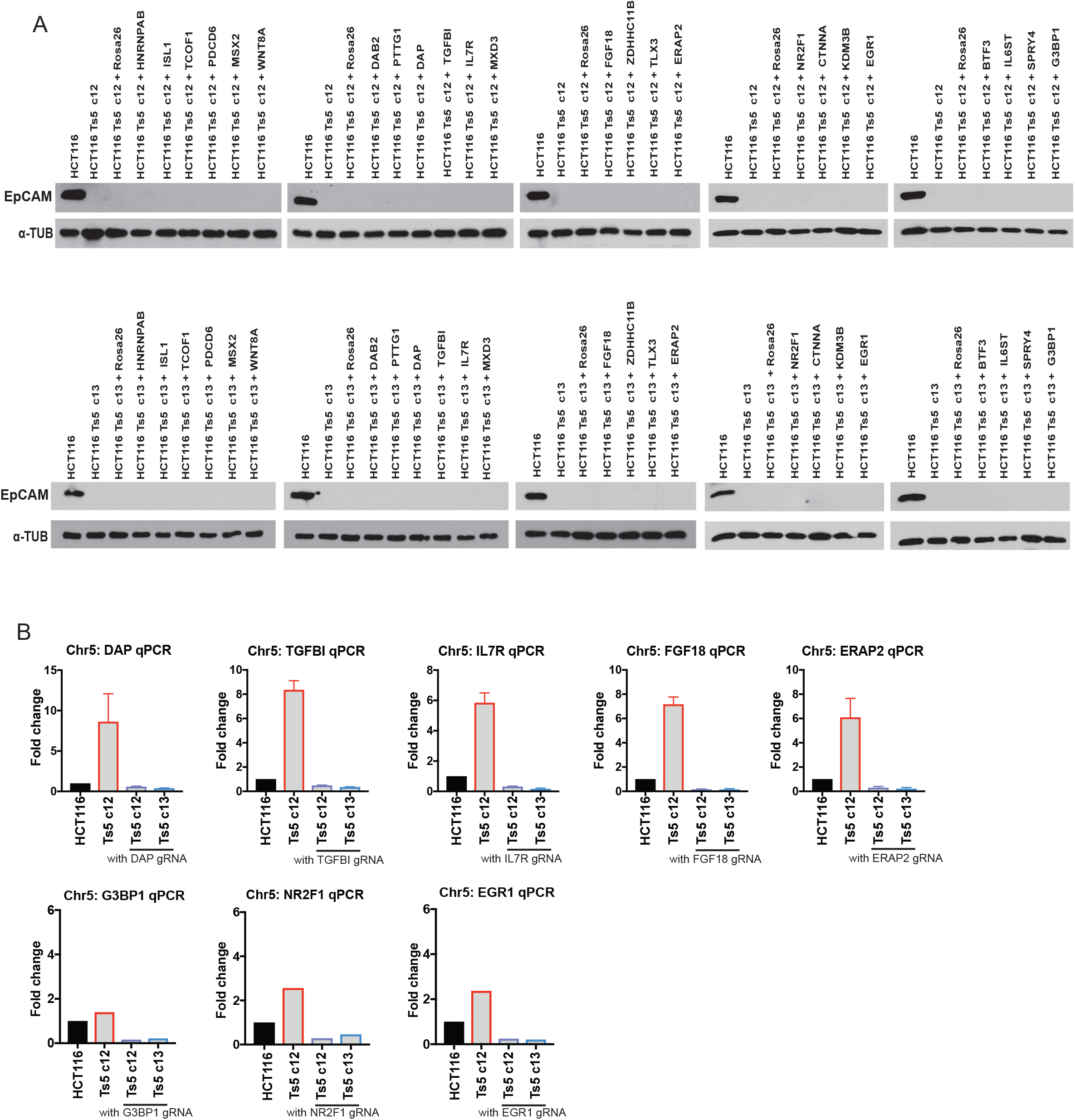
Downregulation of several genes on chromosome 5 does not affect EpCAM silencing. (A) Western blot analysis of EpCAM expression In HCT116 Ts5 clone 12 and clone 13, transduced with 24 gRNAs targeting genes on chromosome 5. (B) qPCR validation of effective CRISPRi knockdown for eight of the guide RNAs used in (A).

**Figure S13.**
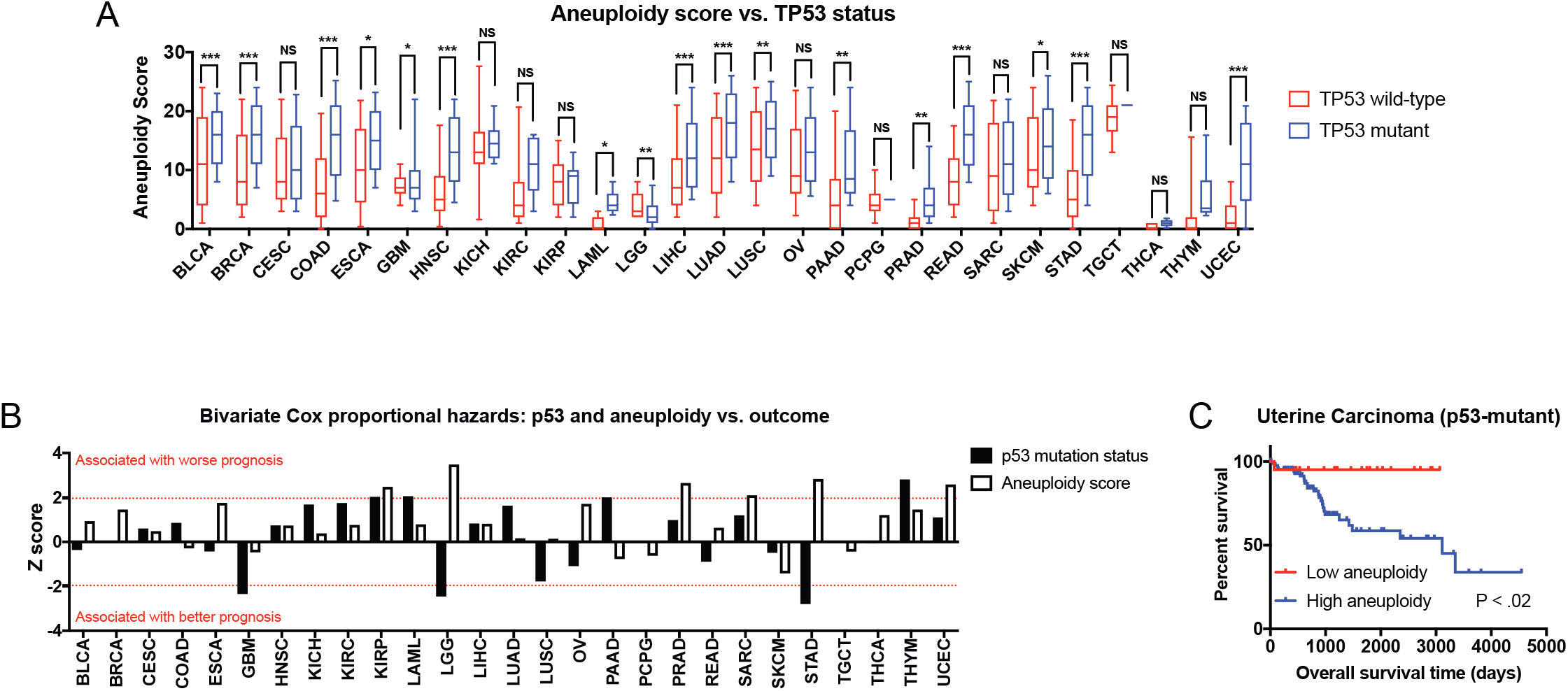
Aneuploidy is a p53-independent prognostic factor in multiple cancer types. (A) The total number of arm-length aneuploidies per sample is plotted for p53-wildtype and p53-mutant tumors from each TCGA cohort. Boxes represent the second and third quartiles, while bars indicate the 10^th^ and 90^th^ percentiles. (B) Z scores from bivariate Cox proportional hazards modeling that combine p53-mutation status and aneuploidy score (split at the 20^th^ and 80^th^ percentiles) are displayed. (C) Kaplan-Meier curve in uterine carcinoma demonstrating that aneuploidy burden is still associated with poor prognosis in p53-mutant tumors.

**Figure S14.**
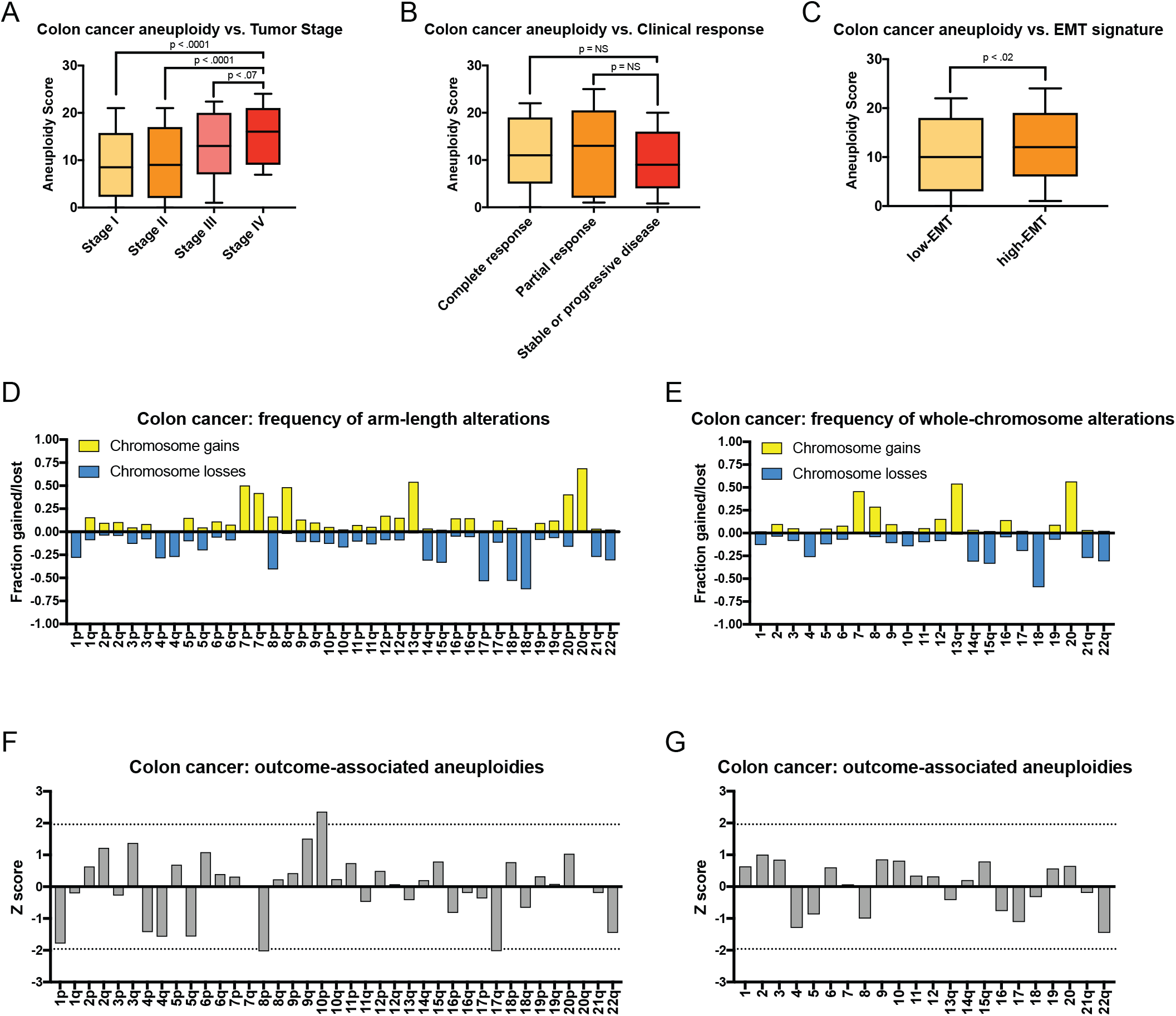
The clinical correlates of colon cancer aneuploidy. (A) Aneuploidy scores by colon cancer stage are displayed. Colon cancer stage information was acquired from (Liu et al. 2018). Boxes represent the second and third quartiles, while bars indicate the 10^th^ and 90^th^ percentiles. (B) Aneuploidy scores by colon cancer treatment response are displayed. Treatment response information was acquired from (Liu et al. 2018). Boxes represent the second and third quartiles, while bars indicate the 10^th^ and 90^th^ percentiles. (C) Colon tumors were split based on the expression of an EMT-related gene signature acquired from (Gibbons and Creighton 2018). “Low-EMT” and “High-EMT” tumors were determined based on the mean expression of the gene signature. Boxes represent the second and third quartiles, while bars indicate the 10^th^ and 90^th^ percentiles. (D) The fraction of tumors that have gained or lost the indicated chromosome is displayed. (E) The fraction of tumors that have gained or lost the indicated chromosome arm is displayed. (F) Z scores for the indicated whole-chromosome aneuploidies are displayed. (G) Z scores for the indicated arm-length aneuploidies are displayed. The dotted lines at Z = 1.96 and −1.96 correspond to P < .05.

**Figure S15.**
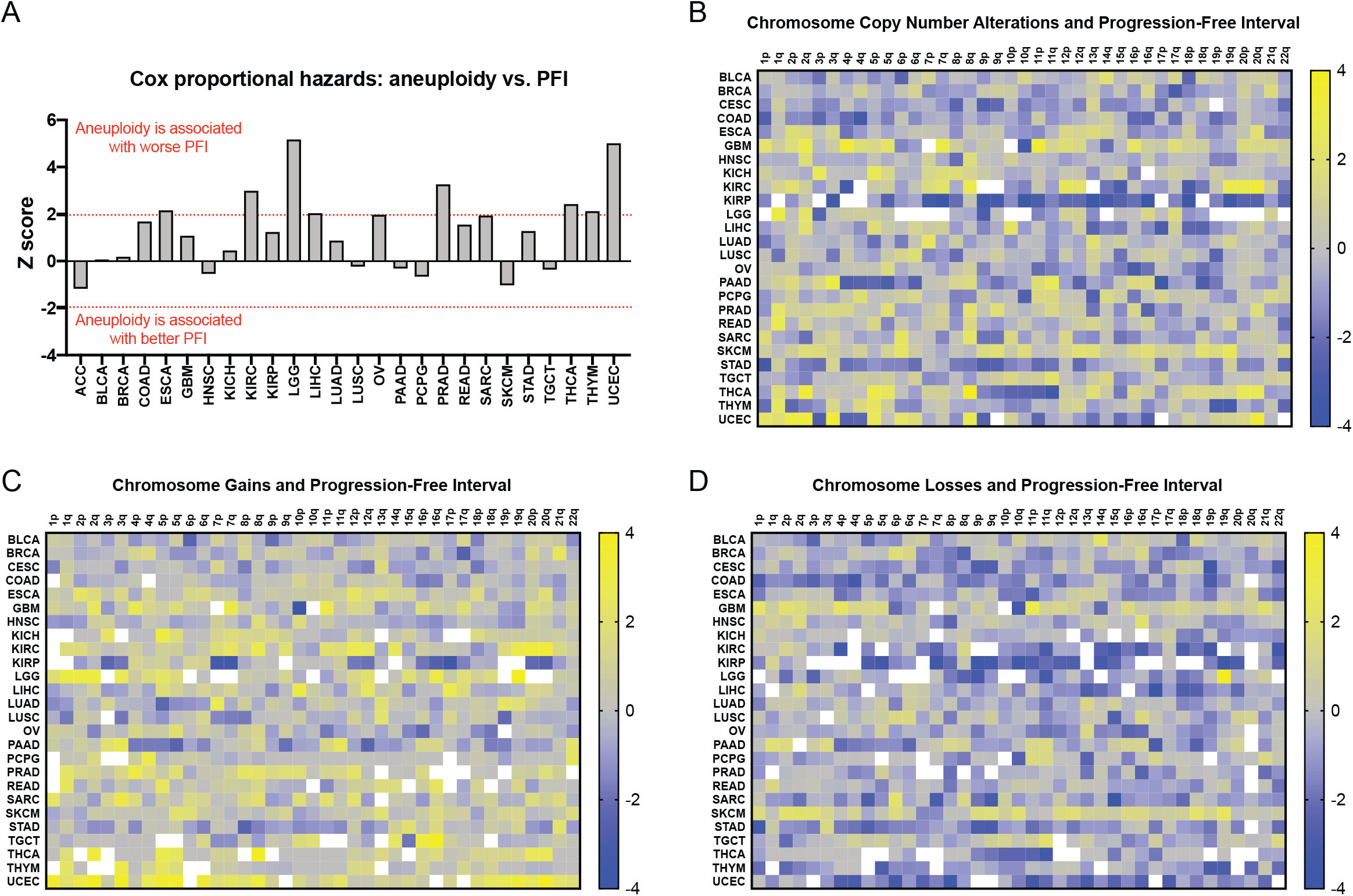
Progression-free interval and cancer aneuploidy. (A) Z scores from Cox-proportional hazards analysis using PFI as an endpoint are displayed. Note that Z > 1.96 indicates a significant association between higher aneuploidy and shorter PFI, while Z < −1.96 indicates a significant association between higher aneuploidy and longer PFI. (B) A heatmap comparing chromosome arm copy number vs. PFI across patients with 26 different types of cancer. The color-bar indicates the Z score from the Cox analysis. For visualization purposes, Z scores were capped at 4 and −4. (C) A heatmap comparing dichotomized chromosome arm copy number vs. patient outcome across patients with 26 different types of cancer. In this analysis, survival was compared between patients in which a given arm was gained and patients in which a given arm was either lost or was copy-neutral. The color-bar indicates the Z score from the Cox analysis. For visualization purposes, Z scores were capped at 4 and −4. (D) A heatmap comparing dichotomized chromosome arm copy number vs. patient outcome across patients with 26 different types of cancer. In this analysis, survival was compared between patients in which a given arm was lost and patients in which a given arm was either gained or was copy-neutral. The color-bar indicates the Z score from the Cox analysis. For visualization purposes, Z scores were capped at 4 and −4.

## References

Andrews P. 2017. MUMdex Software. MUMdex Genome Alignment Anal Softw. https://mumdex.com/ (Accessed February 27, 2019).

Andrews PA, Iossifov I, Kendall J, Marks S, Muthuswamy L, Wang Z, Levy D, Wigler M. 2016. MUMdex: MUM-based structural variation detection. bioRxiv 078261.

Bailey MH, Tokheim C, Porta-Pardo E, Sengupta S, Bertrand D, Weerasinghe A, Colaprico A, Wendl MC, Kim J, Reardon B, et al. 2018. Comprehensive Characterization of Cancer Driver Genes and Mutations. Cell 173: 371–385.e18.

Bakhoum SF, Ngo B, Laughney AM, Cavallo J-A, Murphy CJ, Ly P, Shah P, Sriram RK, Watkins TBK, Taunk NK, et al. 2018. Chromosomal instability drives metastasis through a cytosolic DNA response. Nature 553: 467–472.

Baslan T, Kendall J, Rodgers L, Cox H, Riggs M, Stepansky A, Troge J, Ravi K, Esposito D, Lakshmi B, et al. 2012. Genome-wide copy number analysis of single cells. Nat Protoc 7: 1024–1041.

Becker L, Mito T, Takashima S, Onodera K. 1991. Growth and development of the brain in Down syndrome. Prog Clin Biol Res 373: 133–152.

Carter SL, Cibulskis K, Helman E, McKenna A, Shen H, Zack T, Laird PW, Onofrio RC, Winckler W, Weir BA, et al. 2012. Absolute quantification of somatic DNA alterations in human cancer. Nat Biotechnol 30: 413–421.

Chang K, Marran K, Valentine A, Hannon GJ. 2013. Packaging shRNA retroviruses. Cold Spring Harb Protoc 2013: 734–737.

Chen Y, Li S, Li W, Yang R, Zhang X, Ye Y, Yu J, Ye L, Tang W. 2019. Circulating tumor cells undergoing EMT are poorly correlated with clinical stages or predictive of recurrence in hepatocellular carcinoma. Sci Rep 9: 7084.

Choma D, Daurès J-P, Quantin X, Pujol JL. 2001. Aneuploidy and prognosis of non-small-cell lung cancer: a meta-analysis of published data. Br J Cancer 85: 14–22.

Chunduri NK, Storchová Z. 2019. The diverse consequences of aneuploidy. Nat Cell Biol 21: 54–62.

Collins TJ. 2007. ImageJ for microscopy. BioTechniques 43: S25–S30.

Davoli T, Xu AW, Mengwasser KE, Sack LM, Yoon JC, Park PJ, Elledge SJ. 2013. Cumulative Haploinsufficiency and Triplosensitivity Drive Aneuploidy Patterns to Shape the Cancer Genome. Cell 155: 948–962.

Di Nicolantonio F, Arena S, Gallicchio M, Zecchin D, Martini M, Flonta SE, Stella GM, Lamba S, Cancelliere C, Russo M, et al. 2008. Replacement of normal with mutant alleles in the genome of normal human cells unveils mutation-specific drug responses. Proc Natl Acad Sci U S A 105: 20864–20869.

Domingues PH, Nanduri LSY, Seget K, Venkateswaran SV, Agorku D, Viganó C, Schubert C von, Nigg EA, Swanton C, Sotillo R, et al. 2017. Cellular Prion Protein PrPC and Ecto-5′-Nucleotidase Are Markers of the Cellular Stress Response to Aneuploidy. Cancer Res 77: 2914–2926.

Dongre A, Weinberg RA. 2019. New insights into the mechanisms of epithelial–mesenchymal transition and implications for cancer. Nat Rev Mol Cell Biol 20: 69.

Donnelly N, Passerini V, Dürrbaum M, Stingele S, Storchová Z. 2014. HSF1 deficiency and impaired HSP90-dependent protein folding are hallmarks of aneuploid human cells. EMBO J 33: 2374–2387.

Duesberg P, Li R, Fabarius A, Hehlmann R. 2006. Aneuploidy and Cancer: From Correlation to Causation. Infect Inflamm Impacts Oncog 13: 16–44.

Dürrbaum M, Kruse C, Nieken KJ, Habermann B, Storchová Z. 2018. The deregulated microRNAome contributes to the cellular response to aneuploidy. BMC Genomics 19: 197.

Dürrbaum M, Kuznetsova AY, Passerini V, Stingele S, Stoehr G, Storchová Z. 2014. Unique features of the transcriptional response to model aneuploidy in human cells. BMC Genomics 15: 139.

Fischer KR, Durrans A, Lee S, Sheng J, Li F, Wong STC, Choi H, El Rayes T, Ryu S, Troeger J, et al. 2015. Epithelial-to-mesenchymal transition is not required for lung metastasis but contributes to chemoresistance. Nature 527: 472–476.

Frankfurt OS, Chin JL, Englander LS, Greco WR, Pontes JE, Rustum YM. 1985. Relationship between DNA Ploidy, Glandular Differentiation, and Tumor Spread in Human Prostate Cancer. Cancer Res 45: 1418–1423.

Friedlander ML, Hedley DW, Taylor IW. 1984. Clinical and biological significance of aneuploidy in human tumours. J Clin Pathol 37: 961–974.

Gialeli C, Theocharis AD, Karamanos NK. 2011. Roles of matrix metalloproteinases in cancer progression and their pharmacological targeting. FEBS J 278: 16–27.

Gibbons DL, Creighton CJ. 2018. Pan-cancer survey of epithelial–mesenchymal transition markers across The Cancer Genome Atlas. Dev Dyn Off Publ Am Assoc Anat 247: 555–564.

Giuliano CJ, Lin A, Sheltzer J. 2019. Generating single cell-derived knockout clones in mammalian cells with CRISPR/Cas9. PeerJ Inc. https://peerj.com/preprints/27511 (Accessed February 15, 2019).

Giuliano CJ, Lin A, Smith JC, Palladino AC, Sheltzer JM. 2018. MELK expression correlates with tumor mitotic activity but is not required for cancer growth. eLife 7: e32838.

Glaab WE, Tindall KR. 1997. Mutation rate at the hprt locus in human cancer cell lines with specific mismatch repair-gene defects. Carcinogenesis 18: 1–8.

Gurden MD, Westwood IM, Faisal A, Naud S, Cheung K-MJ, McAndrew C, Wood A, Schmitt J, Boxall K, Mak G, et al. 2015. Naturally Occurring Mutations in the MPS1 Gene Predispose Cells to Kinase Inhibitor Drug Resistance. Cancer Res 75: 3340–3354.

Heerboth S, Housman G, Leary M, Longacre M, Byler S, Lapinska K, Willbanks A, Sarkar S. 2015. EMT and tumor metastasis. Clin Transl Med 4: 6.

Holland AJ, Cleveland DW. 2009. Boveri revisited: chromosomal instability, aneuploidy and tumorigenesis. Nat Rev Mol Cell Biol 10: 478–487.

Horlbeck MA, Gilbert LA, Villalta JE, Adamson B, Pak RA, Chen Y, Fields AP, Park CY, Corn JE, Kampmann M, et al. 2016. Compact and highly active next-generation libraries for CRISPR-mediated gene repression and activation. eLife 5: e19760.

Hume RF, Drugan A, Reichler A, Lampinen J, Martin LS, Johnson MP, Evans MI. 1996. Aneuploidy among prenatally detected neural tube defects. Am J Med Genet 61: 171–173.

Jolly MK, Ware KE, Gilja S, Somarelli JA, Levine H. 2017. EMT and MET: necessary or permissive for metastasis? Mol Oncol 11: 755–769.

Kallioniemi O-P, Hietanen T, Mattila J, Lehtinen M, Lauslahti K, Koivula T. 1987. Aneuploid DNA content and high S-phase fraction of tumour cells are related to poor prognosis in patients with primary breast cancer. Eur J Cancer Clin Oncol 23: 277–282.

Kheir SM, Bines SD, Vonroenn JH, Soong SJ, Urist MM, Coon JS. 1988. Prognostic significance of DNA aneuploidy in stage I cutaneous melanoma. Ann Surg 207: 455–461.

Klaeger S, Heinzlmeir S, Wilhelm M, Polzer H, Vick B, Koenig P-A, Reinecke M, Ruprecht B, Petzoldt S, Meng C, et al. 2017. The target landscape of clinical kinase drugs. Science 358: eaan4368.

Knouse KA, Davoli T, Elledge SJ, Amon A. 2017. Aneuploidy in Cancer: Seq-ing Answers to Old Questions. Annu Rev Cancer Biol 1: 335–354.

Lara-Gonzalez P, Westhorpe FG, Taylor SS. 2012. The Spindle Assembly Checkpoint. Curr Biol 22: R966–R980.

Liu J, Lichtenberg T, Hoadley KA, Poisson LM, Lazar AJ, Cherniack AD, Kovatich AJ, Benz CC, Levine DA, Lee AV, et al. 2018. An Integrated TCGA Pan-Cancer Clinical Data Resource to Drive High-Quality Survival Outcome Analytics. Cell 173: 400–416.e11.

Liu X, Li J, Cadilha BL, Markota A, Voigt C, Huang Z, Lin PP, Wang DD, Dai J, Kranz G, et al. 2019. Epithelial-type systemic breast carcinoma cells with a restricted mesenchymal transition are a major source of metastasis. Sci Adv 5: eaav4275.

López-García C, Sansregret L, Domingo E, McGranahan N, Hobor S, Birkbak NJ, Horswell S, Grönroos E, Favero F, Rowan AJ, et al. 2017. BCL9L Dysfunction Impairs Caspase-2 Expression Permitting Aneuploidy Tolerance in Colorectal Cancer. Cancer Cell 31: 79–93.

Maciejowski J, George KA, Terret M-E, Zhang C, Shokat KM, Jallepalli PV. 2010. Mps1 directs the assembly of Cdc20 inhibitory complexes during interphase and mitosis to control M phase timing and spindle checkpoint signaling. J Cell Biol 190: 89–100.

Mehlen P, Puisieux A. 2006. Metastasis: a question of life or death. Nat Rev Cancer 6: 449–458.

Merkel DE, McGuire WL. 1990. Ploidy, proliferative activity and prognosis. DNA flow cytometry of solid tumors. Cancer 65: 1194–1205.

Morikawa K, Walker SM, Nakajima M, Pathak S, Jessup JM, Fidler IJ. 1988. Influence of Organ Environment on the Growth, Selection, and Metastasis of Human Colon Carcinoma Cells in Nude Mice. Cancer Res 48: 6863–6871.

Pachis ST, Kops GJPL. 2018. Leader of the SAC: molecular mechanisms of Mps1/TTK regulation in mitosis. Open Biol 8. https://www.ncbi.nlm.nih.gov/pmc/articles/PMC6119859/ (Accessed January 22, 2019).

Padmanaban V, Krol I, Suhail Y, Szczerba BM, Aceto N, Bader JS, Ewald AJ. 2019. E-cadherin is required for metastasis in multiple models of breast cancer. Nature 573: 439–444.

Pantel K, Brakenhoff RH. 2004. Dissecting the metastatic cascade. Nat Rev Cancer 4: 448–456.

Passerini V, Ozeri-Galai E, de Pagter MS, Donnelly N, Schmalbrock S, Kloosterman WP, Kerem B, Storchová Z. 2016. The presence of extra chromosomes leads to genomic instability. Nat Commun 7: 10754.

Pastushenko I, Blanpain C. 2019. EMT Transition States during Tumor Progression and Metastasis. Trends Cell Biol 29: 212–226.

Pavelka N, Rancati G, Zhu J, Bradford WD, Saraf A, Florens L, Sanderson BW, Hattem GL, Li R. 2010. Aneuploidy confers quantitative proteome changes and phenotypic variation in budding yeast. Nature 468: 321–325.

Rutledge SD, Douglas TA, Nicholson JM, Vila-Casadesús M, Kantzler CL, Wangsa D, Barroso-Vilares M, Kale SD, Logarinho E, Cimini D. 2016. Selective advantage of trisomic human cells cultured in non-standard conditions. Sci Rep 6: 22828.

Santaguida S, Tighe A, D’Alise AM, Taylor SS, Musacchio A. 2010. Dissecting the role of MPS1 in chromosome biorientation and the spindle checkpoint through the small molecule inhibitor reversine. J Cell Biol 190: 73–87.

Santaguida S, Vasile E, White E, Amon A. 2015. Aneuploidy-induced cellular stresses limit autophagic degradation. Genes Dev. http://genesdev.cshlp.org/content/early/2015/09/22/gad.269118.115 (Accessed September 10, 2016).

Sheltzer JM. 2013. A transcriptional and metabolic signature of primary aneuploidy is present in chromosomally-unstable cancer cells and informs clinical prognosis. Cancer Res 73: 6401–6412.

Sheltzer JM, Amon A. 2011. The aneuploidy paradox: costs and benefits of an incorrect karyotype. Trends Genet TIG 27: 446–453.

Sheltzer JM, Blank HM, Pfau SJ, Tange Y, George BM, Humpton TJ, Brito IL, Hiraoka Y, Niwa O, Amon A. 2011. Aneuploidy Drives Genomic Instability in Yeast. Science 333: 1026–1030.

Sheltzer JM, Ko JH, Replogle JM, Habibe Burgos NC, Chung ES, Meehl CM, Sayles NM, Passerini V, Storchova Z, Amon A. 2017. Single-chromosome Gains Commonly Function as Tumor Suppressors. Cancer Cell 31: 240–255.

Sheltzer JM, Torres EM, Dunham MJ, Amon A. 2012. Transcriptional consequences of aneuploidy. Proc Natl Acad Sci 109: 12644–12649.

Smith JC, Sheltzer JM. 2018. Systematic identification of mutations and copy number alterations associated with cancer patient prognosis ed. J. Settleman. eLife 7: e39217.

Soto M, García-Santisteban I, Krenning L, Medema RH, Raaijmakers JA. 2018. Chromosomes trapped in micronuclei are liable to segregation errors. J Cell Sci 131. https://www.ncbi.nlm.nih.gov/pmc/articles/PMC6051344/ (Accessed October 8, 2019).

Stingele S, Stoehr G, Peplowska K, Cox J, Mann M, Storchova Z. 2012. Global analysis of genome, transcriptome and proteome reveals the response to aneuploidy in human cells. Mol Syst Biol 8: 608.

Taube JH, Herschkowitz JI, Komurov K, Zhou AY, Gupta S, Yang J, Hartwell K, Onder TT, Gupta PB, Evans KW, et al. 2010. Core epithelial-to-mesenchymal transition interactome gene-expression signature is associated with claudin-low and metaplastic breast cancer subtypes. Proc Natl Acad Sci 107: 15449–15454.

Taylor AM, Shih J, Ha G, Gao GF, Zhang X, Berger AC, Schumacher SE, Wang C, Hu H, Liu J, et al. 2018. Genomic and Functional Approaches to Understanding Cancer Aneuploidy. Cancer Cell 33: 676–689.e3.

Thompson SL, Compton DA. 2010. Proliferation of aneuploid human cells is limited by a p53-dependent mechanism. J Cell Biol 188: 369–381.

Viganó C, von Schubert C, Ahrné E, Schmidt A, Lorber T, Bubendorf L, De Vetter JRF, Zaman GJR, Storchova Z, Nigg EA. 2018. Quantitative proteomic and phosphoproteomic comparison of human colon cancer DLD-1 cells differing in ploidy and chromosome stability. Mol Biol Cell 29: 1031–1047.

Wang Z, Andrews P, Kendall J, Ma B, Hakker I, Rodgers L, Ronemus M, Wigler M, Levy D. 2016. SMASH, a fragmentation and sequencing method for genomic copy number analysis. Genome Res 26: 844–851.

Williams BR, Prabhu VR, Hunter KE, Glazier CM, Whittaker CA, Housman DE, Amon A. 2008. Aneuploidy affects proliferation and spontaneous immortalization in mammalian cells. Science 322: 703–709.

Xie Y, Wang A, Lin J, Wu L, Zhang H, Yang X, Wan X, Miao R, Sang X, Zhao H. 2017. Mps1/TTK: a novel target and biomarker for cancer. J Drug Target 25: 112–118.

Zheng X, Carstens JL, Kim J, Scheible M, Kaye J, Sugimoto H, Wu C-C, LeBleu VS, Kalluri R. 2015. Epithelial-to-mesenchymal transition is dispensable for metastasis but induces chemoresistance in pancreatic cancer. Nature 527: 525–530.

